# Spatially Resolved *in vivo* CRISPR Screen Sequencing via Perturb-DBiT

**DOI:** 10.1101/2024.11.18.624106

**Authors:** Alev Baysoy, Xiaolong Tian, Feifei Zhang, Paul Renauer, Zhiliang Bai, Hao Shi, Haikuo Li, Bo Tao, Mingyu Yang, Archibald Enninful, Fu Gao, Guangchuan Wang, Wanqiu Zhang, Thao Tran, Nathan Heath Patterson, Shuozhen Bao, Chuanpeng Dong, Shan Xin, Mei Zhong, Sherri Rankin, Cliff Guy, Yan Wang, Jon P. Connelly, Shondra M. Pruett-Miller, Hongbo Chi, Sidi Chen, Rong Fan

**Affiliations:** Department of Biomedical Engineering, Yale University, New Haven, CT, USA; Department of Genetics, Yale University School of Medicine, New Haven, CT, USA; Department of Immunology, St. Jude Children’s Research Hospital, Memphis, TN, USA; Aspect Analytics NV, Genk, Belgium; Center for Advanced Genome Engineering, St. Jude Children’s Research Hospital, Memphis, TN, USA; Department of Pathology, Yale University School of Medicine, New Haven, CT, USA; Systems Biology Institute, Integrated Science & Technology Center, West Haven, CT, USA; Yale Stem Cell Center and Yale Cancer Center, Yale University School of Medicine, New Haven, CT 06520, USA; Department of Cell Biology, Yale Stem Cell Center, Yale School of Medicine, New Haven, CT, 06520, USA; Human and Translational Immunology, Yale University School of Medicine, New Haven, CT 06520, USA

**Keywords:** Spatial omics, in vivo CRISPR screen, genetic perturbation, whole transcriptome, genome-wide CRIPR library, tumor colonialization, clonal dynamics, tumor immune microenvironment

## Abstract

Perturb-seq enabled the profiling of transcriptional effects of genetic perturbations in single cells but lacks the ability to examine the impact on tissue environments. We present Perturb-DBiT for simultaneous co-sequencing of spatial transcriptome and guide RNAs (gRNAs) on the same tissue section for in vivo CRISPR screen with genome-scale gRNA libraries, offering a comprehensive understanding of how genetic modifications affect cellular behavior and tissue architecture. This platform supports a variety of delivery vectors, gRNA library sizes, and tissue preparations, along with two distinct gRNA capture methods, making it adaptable to a wide range of experimental setups. In applying Perturb-DBiT, we conducted un-biased knockouts of tens of genes or at genome-wide scale across three cancer models. We mapped all gRNAs in individual colonies and corresponding transcriptomes in a human cancer metastatic colonization model, revealing clonal dynamics and cooperation. We also examined the effect of genetic perturbation on the tumor immune microenvironment in an immune-competent syngeneic model, uncovering differential and synergistic perturbations in promoting immune infiltration or suppression in tumors. Perturb-DBiT allows for simultaneously evaluating the impact of each knockout on tumor initiation, development, metastasis, histopathology, and immune landscape. Ultimately, it not only broadens the scope of genetic inquiry, but also lays the groundwork for developing targeted therapeutic strategies.

## INTRODUCTION

CRISPR-Cas9 has emerged as a powerful tool for precise genome editing, enabling genetic manipulation to study functional genomics in a variety of biological systems, including developmental biology^1^, neuroscience^2^, cancer^3^, and immunology research^4^, where it has been demonstrated to uncover gene functions and elucidate tumor initiation, development, and metastasis mechanisms^5–7^ Further, high-content large-scale CRISPR screens^8^ enable the discovery of key genetic targets and have been instrumental in systematically perturbing genes to and analyzing their effects on cellular phenotypes^9,10^. Recent advancements in single-cell multi-omics technologies and pooled single-cell CRISPR screening have revolutionized our understanding from genomics to functional genomics by elucidating how genetic alterations affect transcriptional outputs, influence cellular heterogeneities, and key regulatory pathways at single-cell level ^11–17^. For example, Perturb-seq combines single cell RNA-seq and CRISPR based perturbations to perform many functional genomic assays in a pool^11^ and discover cell-type-specific transcriptional programs in development or pathogenesis^18^. CRISPR droplet sequencing (CROP-seq) enables pooled CRISPR screens with single-cell transcriptome resolution using droplet microfluidics^15^, taking genetic screening into the single-cell era. It was further expanded to the genome scale to map the genotype-phenotype landscape, predicting the function of thousands of genes at single cell resolution^19^. Deep learning of the single-cell perturbation atlas can further integrate disparate information sources and enhance our understanding of complex, heterogeneous, and dynamics cellular systems^20,21^.

However, a critical gap remains in understanding the role of functional genetic perturbation *in vivo* on spatial organization of complex tissues like tumors. The tumor microenvironment (TME) is a spatially organized ecosystem where cell-to-cell interactions and spatial cues play critical roles in cancer development and progression. Understanding how genetic perturbations influence these interactions is essential for developing effective therapies. Spatial CRISPR genomics developed by Brown et al.^22^ uses a ProCode method to detect a set of gRNAs and correlate them with spatial transcriptome from a serial tissue section, uncovering tumor clone-specific transcriptional outputs. Other technologies, largely based on imaging, emerge to address the limitation in spatial context by combining CRISPR-based genetic perturbations with various spatial assays, uncovering some of the unknown drivers of tumor clonality while obtain some information about the spatial architecture of the sample, revealing how specific gene perturbations and their associated phenotypes influence their neighboring tissue or cellular regions^23–30^. However, these methods are yet to realize joint mapping of perturbations and transcriptome on the same tissue section and often limited by the size and pre- defined nature of the CRISPR libraries used.

To overcome these limitations, we introduce “Perturb-DBiT” (Perturbation compatible Deterministic Barcoding in Tissue), a novel unbiased co-profiling technology that integrates spatial transcriptomics and spatial CRISPR screen readouts amenable for genome-scale libraries (Figure 1A). This approach provides a comprehensive understanding of the spatial functional landscape of TME heterogeneity by simultaneously capturing gene expression profiles and the functional consequences of the genetic perturbations on the perturbed cells and local environments across the entire tissue area. Through the preservation of spatial architecture, we provide a method to elucidate how genetic perturbations influence tumor biology, histopathology, tumor clonal expansion, immune cell composition, and cellular state and dynamics. Our method can be potentially expanded to spatial mapping of genetic variants in other high throughput screen systems^31^ or lineage tracing studies^32^.

**Figure 1:**
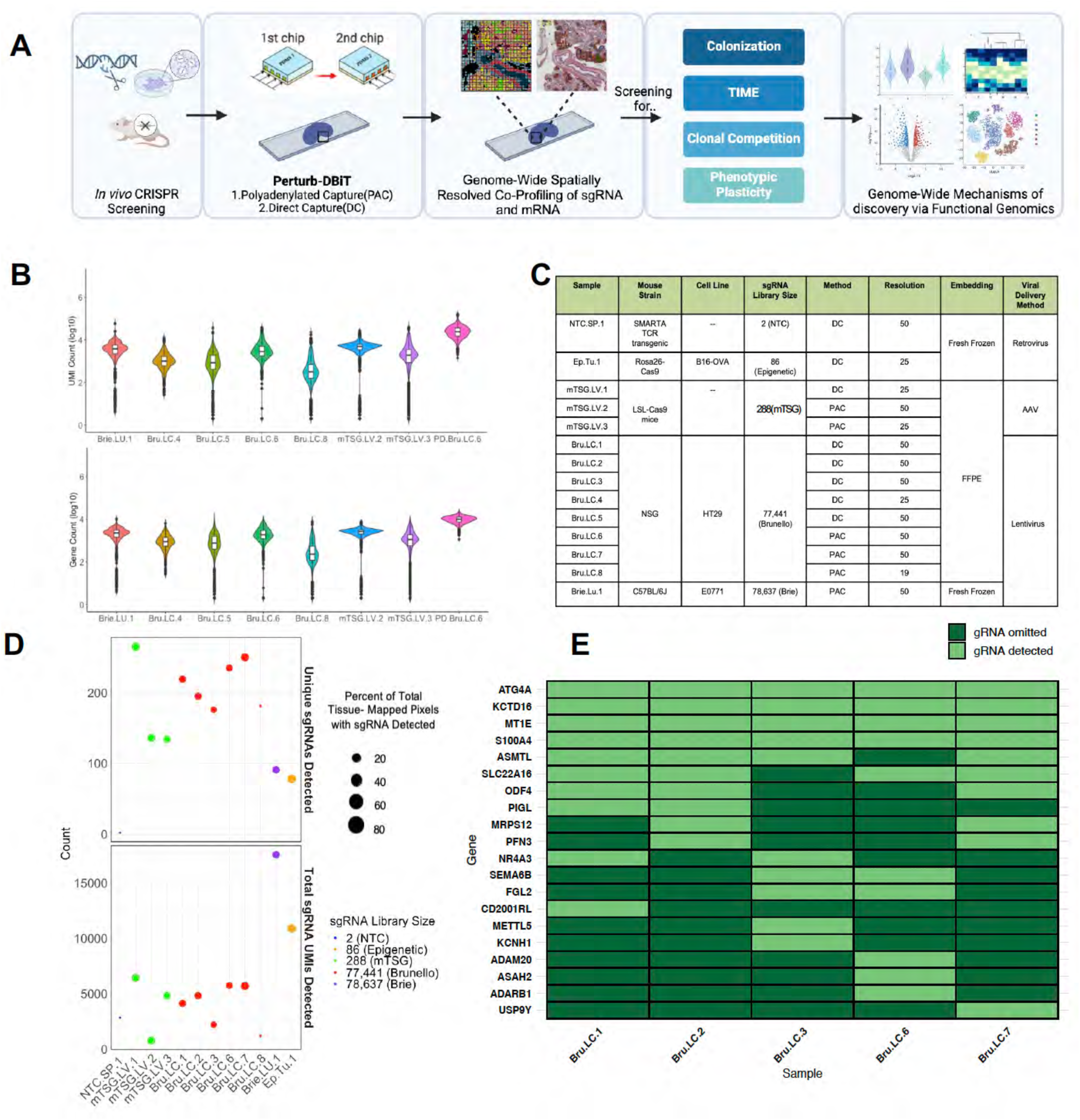
Perturb-DBiT overview and technical performance. (A) Schematic of Perturb-DBiT and its applications. (B) Violin plot of detected gene/UMI counts per spatial pixel from gene expression data across 7 samples utilizing Perturb-DBiT co-profiling. (C) Table of fourteen samples utilizing Perturb-DBiT co-profiling and Perturb-DBiT sgRNA capture in different tissue types, resolutions, viral delivery methods, embeddings, and sgRNA library sizes. (D) Bubble plot depicting Perturb-DBiT sgRNA capture diversity. (E) Table depicting reproducibility of Perturb-DBiT direct capture across five adjacent samples.

## RESULTS

### Perturb-DBiT Workflow

Perturb-DBiT consists of two unique approaches to co-profile single-guide RNA (sgRNA) and transcriptomic mRNA within the same CRISPR-screen-edited tissue, which we demonstrate as polyadenylated capture (PAC) and direct-capture (DC) (Figures S1A, S1B). Both methods are compatible with fresh-frozen (FF) and formalin-fixed paraffin-embedded (FFPE) tissues^33^.

Perturb-DBiT PAC applied to FFPE CRISPR-screen-edited tissue sections initiates with deparaffinization of the tissue section and heat-induced crosslink reversal while FF samples initiate with 1% PFA fixation. Following permeabilization of the tissue, enzyme-induced *in situ* polyadenylation enables the co-detection of both large and small RNA as well as sgRNA by covalently attaching adenosine residues to the 3’ end of RNA molecules, forming a poly(A) tail (Figure S1C). Bulk reverse transcription (RT) is then performed using a biotinylated polydT reverse transcription primer made up of a poly(T) oligo, unique molecular identifier (UMI), and a ligation linker (10mer). Next, spatial barcoding is executed using two polydimethylsiloxane (PDMS) microfluidic chips (or devices), each featuring 50 or 100 parallel microchannels, and each with channels oriented in directions perpendicular to one another^34,35^. Distinct spatial barcodes Ai (i=1-100, 8mer) and Bj (j =1-100, 8mer) are utilized with the first and second PDMS device respectively, with imaging after each barcoding step to preserve the tissue’s region of interest (ROI) for downstream processing. Post-barcoding, the tissue undergoes lysis and template switching followed by PCR to extract barcoded mRNA cDNA and sgRNA for the downstream workflow. Perturb-DBiT PAC PCR utilizes two primers specific to the TSO-end and Barcode B-end of the transcript. Following PCR, left-sided SPRI-select size selection-based separation is then used to achieve separation between the mRNA and sgRNA. As the polyadenylation process adds poly(A) tails to all RNAs, and not just the biomolecules of interest, cDNAs originating from rRNAs were selectively removed from amplicons to circumvent the loss of low-abundance transcripts^36^. After rRNA removal, the separated and purified sgRNA and mRNA cDNA is used as input for library preparation prior to next-generation sequencing.

Perturb-DBiT DC follows similar tissue preparation (Figure S1D). Following tissue permeabilization, bulk reverse transcription is achieved using two distinct RT primers: a biotinylated polyd(T) primer and a direct capture primer. Both primers consist of a UMI and ligation linker 1 or 3, respectively. In addition to these components, the polyd(T) primer uses an oligo dT to capture inherently polyadenylated mRNA while the direct capture primer uses a “capture” sequence that is complementary to the scaffold sequence on the sgRNA constant region. Initially, to optimize this approach, we performed a series of three “bulk” Perturb-DBiT DC tests (forgoing spatial resolution) on adjacent murine spleen sections perturbed with a small 2-non-targeting control (NTC) sgRNA library, utilizing only one barcode *A*_1_, *B*_1_, *C*_1_ and *D*_1_. Each section tested a different ratio of sgRNA to mRNA capture reagents (specific RT primers, barcodes, and primers) (Figure S1E, Table S3). We found that a 1:1 ratio of sgRNA: mRNA capture reagent was most efficient for Perturb-DBiT DC, achieving 87% capture efficiency of both mRNA barcodes A and B and of 1200 sgRNA UMIs with barcode C and D. Utilizing this optimized ratio, after RT, a first round of barcoding is executed using equal proportions of barcode Ai (i=1- 50, 8mer) and barcode Ci (i=1-50, 8mer) where barcode A is compatible with ligation linker 1 (10mer) and barcode C is compatible with ligation linker 3 (10mer). The second round of barcoding is then replicated using barcode Bj (j=1-50, 8mer) and barcode Dj (j=1-50, 8mer). After template switching, and PCR amplification using an additional primer targeting the sgRNA-end of the transcript, a mixture of both sgRNA labeled with spatial barcodes Ci,Dj and mRNA cDNA labeled with spatial barcodes Ai,Bj is collected. Left-sided size selection was used to separate these two products, followed by sgRNA double-sided size-selection. Once correctly sized cDNA and sgRNA counterparts were validated using tapestation or bioanalyzer, these products were used to build libraries prior to next-generation sequencing.

Perturb-DBiT PAC and DC offer two unique methods to co-profile sgRNA and mRNA from CRISPR-screen edited tissue sections while preserving spatial architecture. By combining CRISPR-based genetic perturbations with spatial transcriptomics, Perturb-DBiT provides a powerful yet flexible tool for investigating the spatial organization of gene function and the effects of genetic alterations on cellular behavior, adding a new and integral dimension to spatially resolved multi-omics data (Figures 1A, S1A).

We applied Perturb-DBiT to diverse panel of FFPE and FF samples from various immune-editing models. These samples yielded tissue from SMARTA TCR transgenic spleen, B16-OVA mouse tumor, autochthonous mouse liver tumors, HT29 lung metastatic colonization tissue, and E0771 syngeneic lung colonization tumors, representing a broad spectrum of cancer types and cancer immunology experimental designs (Figure 1C).

Samples were perturbed with sgRNA pooled screens ranging in size from a small pool of 2 or 86 sgRNAs to a larger pool of 288 sgRNAs, and even extending to a genome-wide sgRNA library containing approximately 80,000 gRNAs. Furthermore, we successfully employed Perturb-DBiT with lentivirus, retrovirus, and AAV viral vector delivery methods. This versatility underscores Perturb-DBiT’s adaptability, making it suitable for a broader range of experimental applications. Perturb-DBiT consistently generated high-quality spatial transcriptomics data across diverse tissue types (Figure 1B). Notably, one PAC HT29 lung metastatic colonization FFPE sample yielded 4,220 UMIs and 2,319 unique genes per pixel. On the other hand, an adjacent DC HT29 lung metastatic colonization FFPE sample yielded 1,467 UMIs and 1,130 unique genes per pixel, suggesting that the PAC method may outperform DC in mRNA capture. These results demonstrate Perturb-DBiT’s ability to capture detailed gene expression profiles, even in challenging FFPE samples.

Perturb-DBiT demonstrated robust sgRNA capture performance across a range of CRISPR screen libraries and experimental conditions (Figure 1D, Table S1). We successfully and unbiasedly detected from 2 to 265 unique sgRNAs in our regions of interest (ROI) and between 754 to 17,548 unique total sgRNA UMIs with 1- 106 QC-passed sgRNA UMIs per pixel, dependent on the size of the CRISPR screen library and the sample type. Notably, we were able to detect 5,740 unique sgRNA UMIs (1-26 UMIs/pixel) from a genome-wide- CRISPR screen edited PAC lung metastatic colonization sample, accounting for all 52% of the tissue-containing pixels and representative of 250 unique sgRNAs. To assess the reproducibility of Perturb DBiT’s unbiased sgRNA capture, we analyzed adjacent sections of genome wide CRISPR screen edited HT29 lung metastatic colonization samples with similar ROIs (Figure 1E). Between four adjacent sections, we were consistently able to capture the top 6 sgRNAs hits, demonstrating high reproducibility of Perturb-DBiT in capturing biologically relevant genetic perturbations. With this demonstration, it is important to note that many of the specific genes detected across experiments, even in adjacent tissues, may slightly vary between samples due to subtle spatial variations in gene expression and functional phenotypes. These differences could be attributed to clonal heterogeneity, microenvironmental influences, or stochastic effects of CRISPR-mediated gene editing. Perturb-DBiT is capable of accurately capturing spatial variations in sgRNA abundance within the tissue.

### Perturb-DBiT ensures precise sgRNA fidelity and recapitulation across small and medium sized CRISPR screening libraries

To demonstrate Perturb-DBiT’s ability to profile sgRNA while preserving spatial context, we utilized three immune-editing biological models with small (2 or 86 sgRNA) and medium-sized (288 sgRNA) CRISPR screening libraries and different modes of CRISPR vector delivery (Figure 2A). For the small sgRNA library, B16 (melanoma) OVA-Cas9-GFP cell lines were transduced with retroviral CRISPR sgRNA library targeting 38 epigenetic genes (two sgRNA sequences per gene and 10 NTCs). Here, a modified ametrine-expressing retroviral vector was generated using a dual-guide direct capture lentiviral sgRNA vector. Fourteen days after injection, B16-OVA tumors were harvested and sectioned for H&E staining and Perturb-DBiT-PAC sgRNA profiling. Perturb-DBiT enabled the efficient capture of 78 out of 86 sgRNAs (1-35 UMIs/pixel), with sgRNAs detected in 90% of all tissue-bearing pixels, demonstrating robust spatial coverage. This unique dual-guide library allows for accessing the capture accuracy of Perturb-DBiT as validated by the significant overlap observed between UMIs for sgRNA 1 and sgRNA 2 of each gene target relative to the ROI (Figure 2B). We also applied the same dual-guide direct capture lentiviral sgRNA vector with a 2-NTC library to an immune-editing model transducing only CD4+ SMARTA T cells^37^ in splenic tissue from a TCR transgenic model. Fresh frozen sections were prepared 6 days after CRISPR mutagenesis and Perturb-DBIT DC revealed significant overlap between the two NTC sgRNAs’ UMIs (1-5 sgRNA UMI/pixel) relative to the ROI. This overlap was further validated with the use of IF-stained images to reveal sgRNA alignment with the B-cell zone, suggesting that CD4+ SMARTA cells are mostly differentiated to follicular helper (Tfh) cells and interact with B cells at day 6 post-LCMV infection^38^ (Figure S2A). Together, these unique models demonstrate the feasibility of Perturb-DBiT DC and PAC methods for small-library sgRNA capture and in both tumor and immune cells.

**Figure 2:**
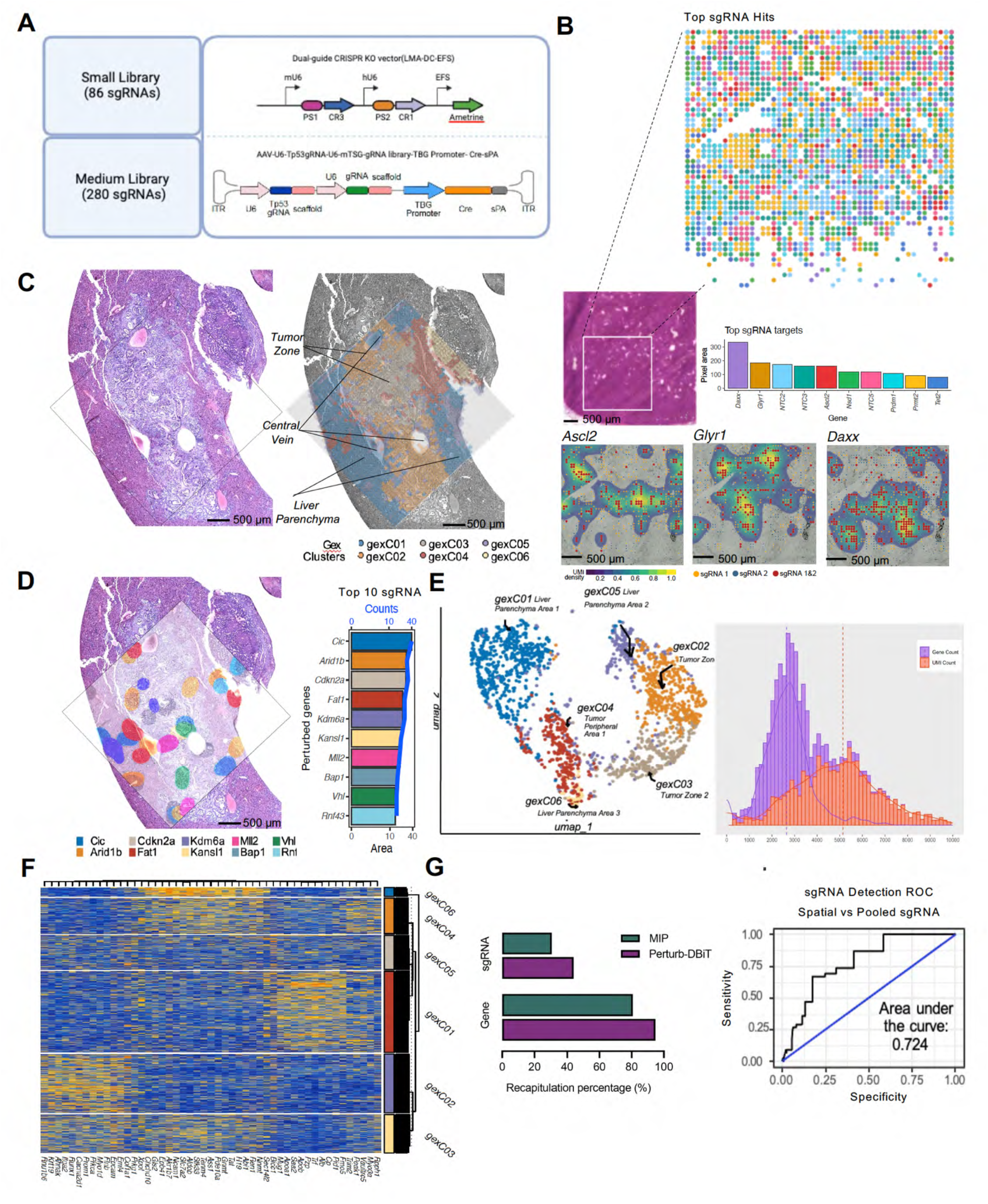
Perturb-DBiT demonstrates robust performance and reveals tumor clonal heterogeneity in small and medium-sized CRISPR-screening libraries. (A) Top: Schematic of the dual sgRNA CRISPR KO retroviral vector used for Perturb-DBiT with the small (2 and 86 sgRNA) libraries. Bottom: Schematic of the AAV-CRISPR vector containing two sgRNA expressing cassettes used for Perturb-DBiT with the medium (288 sgRNA library). (B) Small library guide detection (86 sgRNAs) in Cas9-expressing tumor cells in a tumor tissue using a library of 38 epigenetic regulators and 10 non-targeting controls (NTC). Top: Top sgRNA hit spatial plot and adjacent H&E image corresponding to the region of interest. Bottom: Top sgRNAs are presented with points atop a 2D density map (sgRNA1 and sgRNA2 overlap = blue-to-gold). **(C)** Detection of a medium-sized guide library (288 sgRNAs) in a murine model of autochthonous liver cancer. Left: Liver tissue stained with hematoxylin and eosin (H&E), demonstrating the ROI used for Perturb-DBiT. Right: spatial clustering of gene expression data overlaid on brightfield image, labeled with pathologist annotations. **(D)** Left: Spatial visualization of the top gene perturbations, depicted by color-coded 2D-density maps atop an autochthonous liver tumor ROI. Right: Bar plot of the top 10 gene perturbations detected by 50-micron Perturb- DBiT. The area and total counts of the gene perturbations are presented by bars and blue line, respectively. **(E)** Left: UMAP dimensional reduction embedding of gene expression, leading to distinct spatial clusters. Right: Distribution of detected gene/UMI counts per spatial pixel from gene expression data. (F) Heat map of differentially expressed (DE) genes between GEX pixel clusters. The top 15 expressed DE genes are shown for each cluster with z-score values, arranged by hierarchical clustering of genes, pixels, and clusters. **(G)** Left: Recapitulation analysis comparing sgRNA capture accuracy between Perturb-DBiT and the traditional pooled sequencing method (MIP). Right: Receiver-operator curve comparing the sgRNA detection by 50- micron Perturb-DBiT vs the traditional pooled sequencing method (MIP).

We next extended Perturb-DBiT to FFPE liver primary tumor samples, using a larger CRISPR library (288 sgRNAs) to assess sgRNA capture accuracy. We applied Perturb-DBiT to profile sgRNAs from previously published CRISPR genetically engineered mouse models (CRISPR-GEMMs) of liver cancer (Figure S2C), utilizing a mouse tumor suppressor gene (mTSG) CRISPR library, targeting the 56 most frequently mutated tumor suppressor genes from pan-cancer datasets within The Cancer Genome Atlas (TCGA) ^39,40^. The mTSG RNA library was delivered by AAV into LSL-Cas9 knock-in mice^41^, utilizing an all-in-one construct that included both sgRNA and a liver specific promoter (TBG promoter) to drive Cre specific expression in liver cells. AAV-CRISPR-mediated pooled mutagenesis drives autochthonous liver tumorigenesis in this immunocompetent mouse. Usually, in this autochthonous tumor model, the loss of two TSGs should be sufficient for tumor initiation. Thus, our clever AAV-CRISPR vector was designed with two sgRNA expression cassettes^42^, one targeting the TP53 gene, and the other sgRNA targeting a specific TSG. CRISPR-GEMMs enable generation of genetically complex tumors in individual mice that reflect the heterogeneous composition of human tumors^39,40^. Following mutagenesis, the mouse received immune checkpoint blockade (ICB) treatment of anti-PD-1 (aPD-1) and samples were processed for Perturb-DBiT DC sgRNA capture. SgRNAs were detected in nearly 75% of the ROI profiled, with 1-106 UMIs detected per pixel (Figure S2B). Notably, Perturb-DBiT captured 263 of the 288 sgRNAs in the mTSG panel. Since AAV vectors do not integrate into the host cells’ genomic DNA, unlike a regular lentiviral CRISPR screen, the previous study used molecular inversion probe (MIP) to capture the mutational landscape and test sgRNA activity. Comparatively, the previously published MIP analysis from multiple sections of the same tissue block detected 86 unique sgRNAs overlapping with our data, validating that Perturb-DBiT recapitulates 100% of the enriched sgRNAs from the MIP analysis^7,42^.

Moreover, Perturb-DBiT demonstrates remarkable sensitivity, detecting roughly 3-fold the amount of sgRNAs captured by MIP in just a 1.25x1.25 mm ROI.

### Perturb-DBiT enhanced sgRNA capture and transcriptome spatial mapping reveals tumor origin and oncogenic pathways in liver CRISPR-mediated genetically engineered mouse models (GEMMs)

After confirming successful recapitulation of sgRNAs in the PD-1-treated liver CRISPR-GEMM sample, we applied Perturb-DBiT PAC co-profiling to an adjacent section of the same treated liver block using a 50-micron device to capture a larger ROI (2.5x2.5mm). Perturb-DBiT detected an average of 5,092 UMIs/pixel and 2,595 genes/pixel and unsupervised clustering revealed 6 distinct transcriptomic clusters characterized by differentially expressed gene markers. Further, the spatial clustering aligned closely with the histology from an adjacent section stained with hematoxylin and eosin (H&E), delineating tumor zones, liver parenchyma, and the central vein (Figure 2C, 2E, and 2F). H&E analysis revealed carcinoma-like features, indicative of an epithelial origin for the primary tumor. Consistent with this, the analysis of differentially expressed genes (DEGs) in tumor spatial clusters gexC02 and gexC03 revealed upregulation of *Epcam, Krt19* and *Anhak*, markers commonly associated with epithelial cells. The upregulation of *Prom1* further suggested the presence of stem cell-like properties, consistent with its identification as an important liver cancer stem cell (CSC) and the role of Prom1-derived cells in expanding epithelial tumor cells in human hepatocellular carcinoma (HCC)^43^.

Perturb-DBiT PAC again demonstrated superior sgRNA capture efficiency of the mTSG-targeting CRISPR-screen library compared to traditional pooled sequencing (MIP). In the same tissue section, Perturb-DBiT captured 43% of total sgRNAs, while MIP captured only 30%^42^. Additionally, Perturb-DBiT recapitulated 95% of the 57 mTSG target genes, capturing more sgRNA diversity than MIP (80%) (Figure 2G). Receiver-operator curve analysis revealed a trend toward improved true sgRNA detection accuracy compared to the published MIP data with an AUC of 0.724. While an acceptable model for correct sgRNA classification, this recapitulation result is particularly remarkable given the spatial sequencing nature of Perturb-DBIT and the relatively small ROI analyzed compared to the traditional pooled sequencing. We identified the top 10 sgRNA hits using Perturb-DBiT, which exhibited a heterogeneous spatial distribution, reflecting the inherent genetic and phenotypic diversity of primary tumors (Figure 2D). We next sought to integrate findings from sgRNA and transcriptomics profiling through use of dimensional reduction of Mixscape perturbation scores (pUMAP), which allowed us to visualize and differentiate cell populations based on their response to specific perturbations (Figure S2D). This approach revealed distinct clusters defined by their major perturbation (>=20% of clustered pixels) and included non-perturbed(NP) pixels serving as a control/reference point to enable a direct comparison between perturbed and unperturbed cells. Notably, top pUMAP cluster *Mll2,* also known as *Kmt2d,* also emerged as top depleted hit from the MIP data yet is enriched in this dataset. We utilized violin plots to analyze oncogene expression between the different pUMAP clusters, which correspond to distinct cellular populations or functional states within the tissue (Figure S2D). pUMAP cluster *Cdkn2a*, a well-known tumor suppressor and senescence gene, is found to impact multiple oncogenes in this analysis, suggesting it as an important player in the network of oncogenic regulation. Further, genetic alterations or deletion of *Cdkn2a* are indicative of HCC tumors with poor prognosis, suggesting a role of the P21 pathway in tumor aggressiveness^44^.

Differential expression (DE) analysis of each pUMAP cluster using pseudo-replicate Wilcox tests (Figure S2E) revealed significant insights. The *Bap1* pUMAP cluster, demonstrated upregulation of *Zscan20* – critical for liver carcinoma development – indicating compensatory transcriptional responses to disrupted chromatin dynamics^45^. Gene set enrichment analyses (Figure S2F) further identified key biological pathways, notably in the *Cdkn2a* cluster, which is linked to negative regulation of apoptosis. This suggests that *Cdkn2a* knockout may allow damaged or mutated cells to survive, contributing to cancer progression, a mechanism that has been seen in colorectal cancer^46^. These results underscore the roles of functional genomics in shaping liver tumor biology and offer potential insights for therapeutic interventions. Further, the spatial distribution of sgRNAs and transcriptomic profiles might reflect responses to immune checkpoint inhibition by PD-1, which can alter gene expression patterns and TME architecture.

### Perturb-DBiT reveals clonality within HT29 lung metastatic colonization model using a genome-wide *in vivo* CRISPR library

Given Perturb-DBiT’s demonstrated efficacy in profiling small and medium-sized sgRNA libraries, we sought to expand its application to genome-wide human CRISPR knockout pooled library (Brunello). This library contains 77,441 sgRNAs targeting 19,114 genes, with an additional 1,000 non-targeting controls. The intravenous injection of tumor cells and subsequent lung metastatic colonization model, a well-established platform for evaluating humanized cancer metastasis, was ideal for assessing the role of sgRNA in tumor colonization and metastatic progression. We transduced HT29 cells, a highly metastatic human colon cancer cell line, with a pool of Brunello lentiviral vectors encoding the genome-wide CRISPR-Cas9 gene knockout library, to ensure that each tumor cell was infected by only one sgRNA. Each lentiviral vector contains one sgRNA from the Brunello library (Figure S3A). Intravenous injection of HT29-Cas9-Brunello library cells into 8-week-old NSG mice led to tumor colony formation^47^. After 35 days, when tumor colony size is moderate as detected by IVIS, lung tissues were collected for Perturb-DBiT PAC using a 50-micron device. Within the ROI, we identified 235 unique sgRNAs and a total of 5,740 sgRNA UMIs. To visualize the spatial distribution of these sgRNAs, we mapped all the sgRNA hits onto the H&E image of an adjacent tissue section using a platform provided by Aspect Analytics (Genk, Belgium). Top 10 sgRNA hits were highlighted and as expected, most of these hits were located within the tumor colonies (Figures 3A,3B, and S3B). Interestingly, six of the ten most enriched sgRNAs showed clustering within tumor colonies with instances of colocalization between specific sgRNA pairs, including (*S100A4*/*SLC22A16*; *SEMA6B*/*FGL2*; *MT1E*/*KCTD16*) (Figure 3B). The spatial colocalization of these sgRNAs suggest that they may confer a selective advantage to the tumor cells in which they are expressed and a possible synergistic interaction or clonal cooperation among these genes.

**Figure 3.**
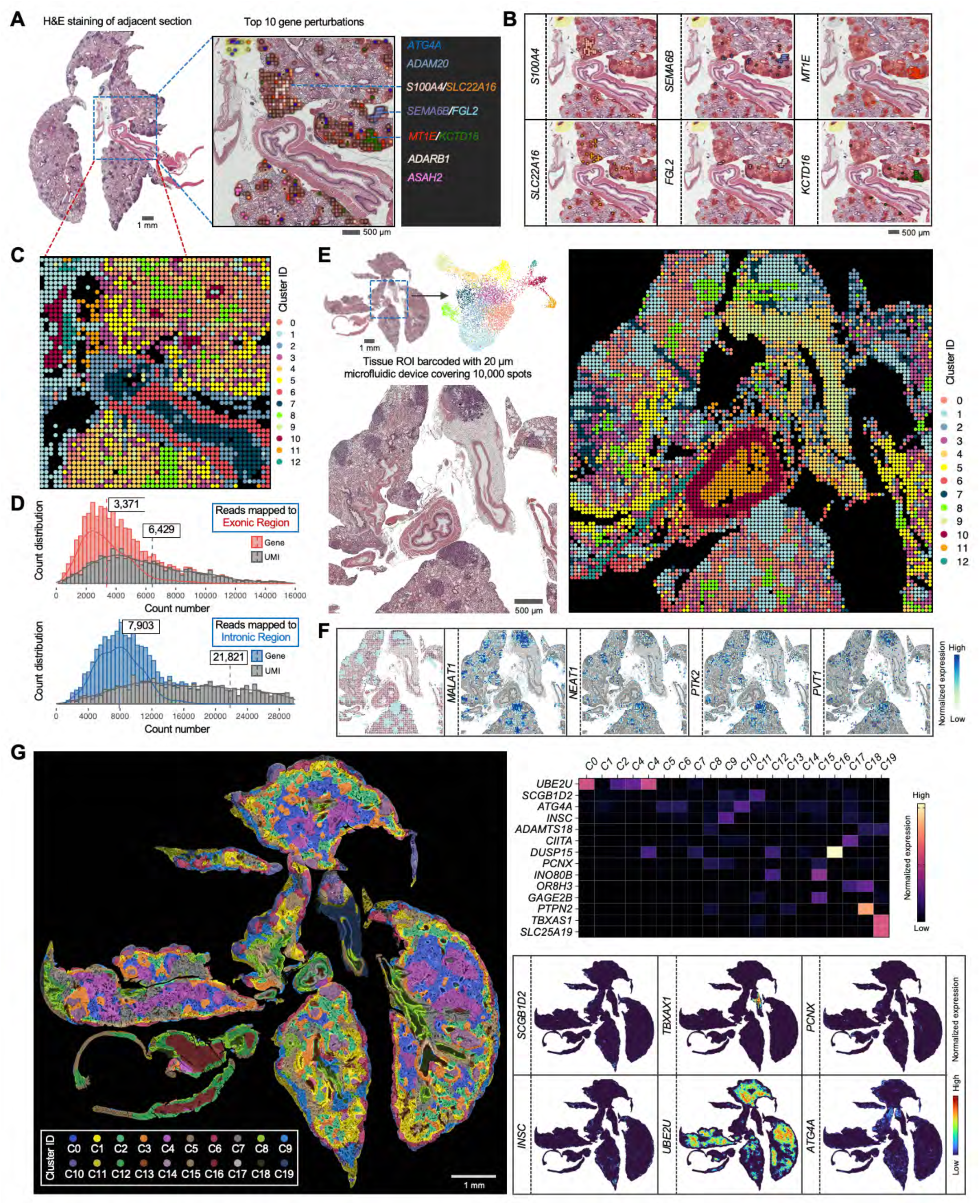
Genome-wide Perturb-DBiT mapping reveals high-resolution tumor clonality by integrating with histology. (A) Spatial mapping of tissue sections collected from HT29 lung metastatic colonization model transduced with genome-wide Brunello sgRNA library. Left: H&E staining of an adjacent section with region of interest (ROI) indicated with blue square. Right: combined spatial distribution of top 10 sgRNAs. (B) Spatial colocalization of top enriched sgRNA pairs. (C) Unsupervised clustering of the combined exonic and intronic expression matrix revealed 20 transcriptomic clusters. (D) Distribution of detected gene/UMI counts per spatial pixel from reads mapped to exonic or intronic region. (E) Spatial mapping of the lung section using 20 μm device covering 100,000 pixels. Left top: H&E of an adjacent section and the UMAP clustering analysis. Left bottom: H&E of the enlarged ROI region. Right: Spatial UMAP showing 13 transcriptomic clusters. (F) Spatial distribution of cluster 1 overlapped with the H&E image, and the top genes defining this cluster. (G) Super-resolved spatial clustering of the top 16 sgRNA profile and representative sgRNA expression enhanced with iStar. These sgRNAs exhibit cluster-specific imputed expression. See also Figures S3.

We next evaluated the gene capture efficiency of Perturb-DBiT on this lung section. Notably, while reads mapped to exonic regions yielded an average of 3,371 genes and 6,429 UMIs per pixel, Perturb-DBiT detected a substantially higher number of intronic molecules, likely attributed to the polyadenylation process, resulting in a mean of 7,903 genes and 21,821 UMIs per pixel (Figure 3D). By combining both exonic and intronic gene data while preserving their distinct identities, unsupervised clustering analysis revealed 13 clusters, each characterized by a unique expression profile (Figures 3C, S3C). The spatial distribution of these clusters closely aligned with the histological structures observed in H&E staining (Figure 3A). To enhance spatial mapping to a near-cellular level while maintaining tissue coverage area, we employed a 100x100 microfluidic device with 20-micron pixel size to spatially barcode 100,000 pixels on an adjacent lung section (Figure 3E).

Clustering analysis of the combined exonic and intronic expression matrix identified 13 clusters that exhibited strong concordance with the histology. The spatial distribution of these clusters revealed distinct and unique patterns (Figure S3D), highlighting Perturb-DBiT’s ability to accurately map fine tissue structures with improved spatial resolution. Particularly, cluster 1 closely mirrored the distribution of tumor regions, as evidenced by its strong alignment with H&E staining patterns of the tumor zones (Figure 3F). Examining the top differentially expressed genes (DEGs) defining this cluster, *MALAT1* emerged as a key regulator of lung cancer metastasis^48^,while *PTK2* was identified as a promoter of metastasis through its role in regulating tumor cell motility and invasion^49^.Additionally, *NEAT1* was linked to tumor cell metabolism^50^,and *PVT1* was found to function as an oncogene involved in the TGF-β signaling pathway^51^. Notably, *MALAT1, NEAT1,* and *PVT1* are non-coding RNAs.

Recognizing the importance of high-resolution histological images for identifying spatial patterns and leveraging recent advances in machine learning for spatial omics data integration, we applied iStar^52^, a computational framework, to fuse Perturb-DBiT data with H&E histology images. Using the top 3,000 DEGs as input, this approach generated a super-resolved, single-cell transcriptomic map, accurately classifying lung tumor cells into clusters G6, G11, and G13 based on their cellular heterogeneity (Figure S3E). Next, we aimed to perform this imputation analysis using the sgRNA expression profile to link the distribution of sgRNAs with tumor colonies. However, the spatial structure of tumor cells was not clearly revealed using the total number of sgRNAs as input (Figure S3F). Given the high accuracy of top enriched sgRNAs in identifying tumor colonies (Figure 3A), we selected 15 sgRNAs—present in the majority of sgRNA-containing pixels (77.6%)—along with a non-targeting control sgRNA, as the input matrix for the iStar analysis. This analysis spatially resolved 20 clusters, with most sgRNAs exhibiting cluster-specific imputed expression (Figure 3G). For instance, cluster C0 showed enriched expression of *UBE2U*, while cluster C16 had notably high expression of *DUSP15*. The distribution of these clusters strongly correlated with tissue structures, with clusters C10 and C14 predominantly overlapping with tumor cell regions. These findings suggest that the combination of Perturb-DBiT and iStar effectively reveals tumor clonal patterns through both gene and top sgRNA expression profiles.

To validate the top sgRNA hits detected by Perturb-DBiT in tumor colony formation, we conducted a clonogenic assay of HT29 cells in vitro (Figure S4A). Briefly, HT29-Cas9 cells were transduced with either a non-targeting control, *AAVS1* sgRNA control, or target sgRNA lentivirus. *AAVS1* is a well-characterized “safe harbor” locus in the human genome, unlikely to disrupt essential genes if edited and is a reliable positive control^53^. After 3-day selection via puromycin and T7E1 assay validation (Figure S4B) of sgRNA genome editing, single tumor cells were seeded at a density of 5,000 cells and resulting cell colonies were analyzed after two weeks. Colony formation assays validated tumor suppressor function of *SEMA6B, MT1E, S100A, and ASAH2* from the Perturb-DBiT top ten enriched hits (Figures 4A, S4C).This was exemplified by the inclusion of four previously reported tumor suppressors (*ARHGAP6, CREBZF, DHRS7C*, and *NR4A3*)^54–57^, whose corresponding sgRNAs were abundantly detected by Perturb-DBiT. Knockout of these genes, except for DHRS7C, which was shown to promote tumor colony growth (Figures S4C, S4D). We next implemented hazard ratios of gene perturbations across various cancer types depicting potential tumor suppressor and promotor activity (Figure 4B). Notably, in kidney renal clear cell carcinoma (KIRC), most of the top ten sgRNA hits are classified as tumor suppressors, suggesting a dominant role for growth inhibition. In contrast, other cancers including HT29, which is classified as colorectal adenocarcinoma (COAD), exhibit a mix of tumor-suppressing and tumor-promoting activities. These results suggest the exceptional performance of Perturb-DBiT in identifying functional perturbations in a spatially relevant context across the genome-wide sgRNA library.

**Figure 4.**
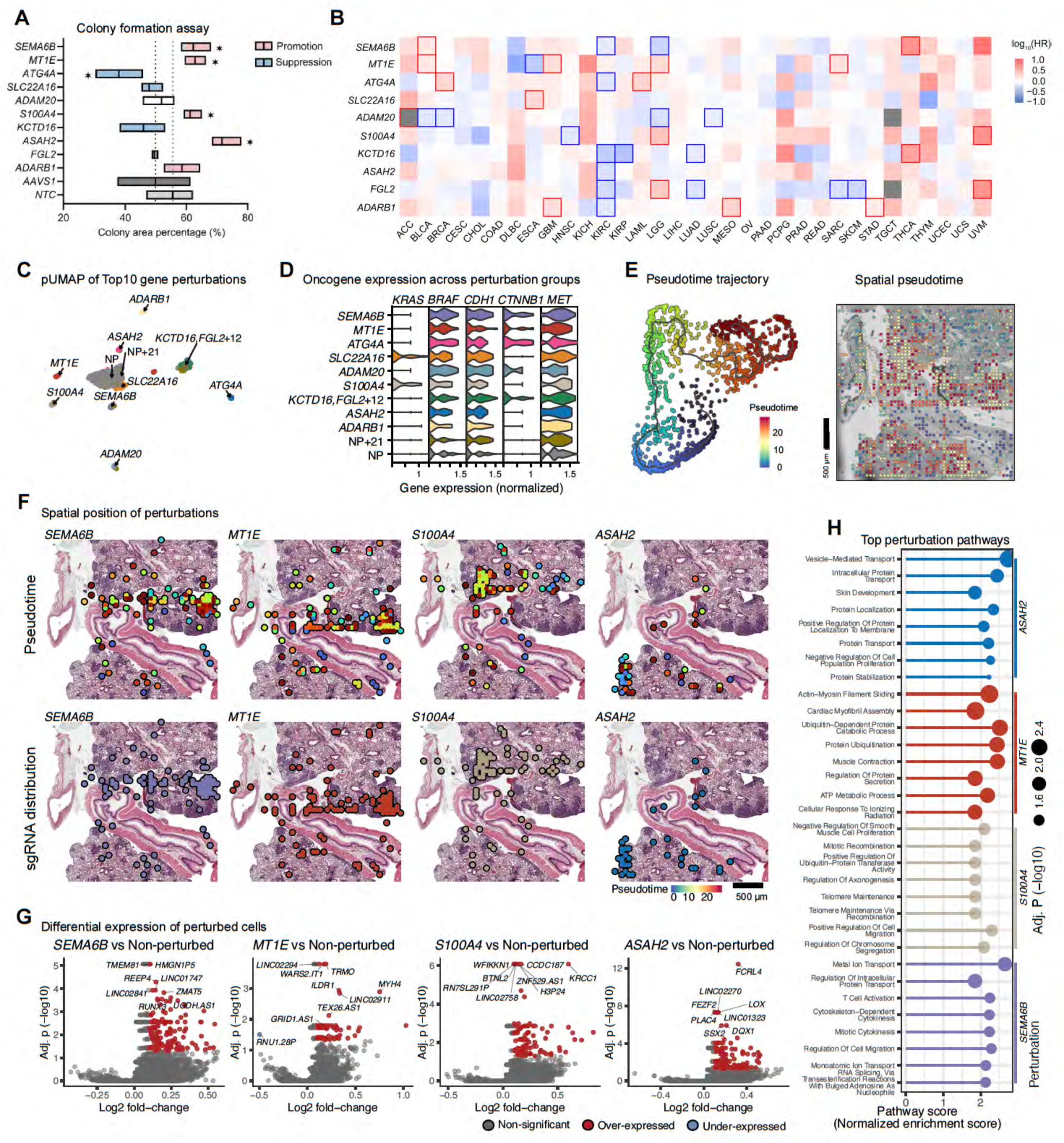
Perturb-DBiT reveals distinct tumor suppressor/promoter programs. (A) Colony formation assay results of top enriched sgRNA from HT29 lung metastatic colonization model. (B) Hazard Ratio plot of top enriched sgRNA from HT29 lung metastatic colonization model. Squares surrounded by colored boxes represent significant differences in hazard ratios. (C) UMAP visualization of pixels based on the dimensional reduction of Mixscape perturbation scores (pUMAP). Non-perturbed (NT) pixels were included in the analysis to serve as a control/reference point. Additionally, pixels were clustered (shown by color and outline) then named by the major perturbations of the cluster (>= 20% of clustered pixels). One cluster with NT and 21 minor perturbations (< 20% of cluster) was named “NT+21”. (D) Violin plot of the expression of 5 selected oncogenes, compared between pUMAP clusters. (E) Monocle3 pseudotime analysis of sgRNA detected from HT29 lung metastatic colonization model. (F) Top: Pseudotime analysis of top 5 enriched sgRNAs detected by Perturb-DBiT. Bottom: spatial mapping of top 5 enriched sgRNAs detected by Perturb-DBiT, overlaid on the brightfield image of the ROI. (G, H) Volcano plots of the transcriptomic DEG analysis for top enriched sgRNA perturbations and the top 8 most significant DE genes were labeled for selected analyses and gene ontology analysis (H).

We next sought to investigate the effect of CRISPR perturbations on spatial transcriptomic profiles through Perturb-DBiT. First, we examined the relationships between these top perturbations and key oncogenic pathways. pUMAP analysis revealed clusters defined by their major perturbation (>=20% of clustered pixels) and included NT pixels serving as a control, highlighting the specific transcriptional and phenotypic changes associated with each sgRNA perturbation, providing deeper insights into how gene knockouts influence cellular states and behaviors (Figure 4C). Further, we compared the effects of each pUMAP cluster on genes associated with critical HT29 tumor biomarkers: *KRAS, BRAF, CDH1, CYNNB1,* and *MET* (Figure 4D). This analysis revealed the diverse functional consequences of the top hits identified by Perturb-DBiT across various oncogenic drivers. Moreover, these results suggest that the top sgRNA perturbations not only vary in their overall effect on cancer types, but also show specificity in how they interact with distinct oncogenic pathways, highlighting Perturb-DBiT’s application to elucidating potential pathway-specific vulnerabilities that could be therapeutically targeted. Secondly, we observed that the spatial distribution of these top sgRNA hits typically follows a pattern of both dispersion and concentration within different tumor colonies (Figure 3B). This suggests that sgRNA enrichment might be indicative of the direction of tumor development. Consequently, by using Monacle3^58^, we conducted pseudotime analysis with spatial transcriptomics data from tumor cells containing sgRNAs. Intriguingly, the distribution patterns of the sgRNAs for *SEMA6B*, *MT1E*, and *S100A4* - three of the four validated tumor suppressors- align well with the tumor pseudotime trajectory, suggesting the formation of diverse tumor genotypes in a single pooled screen that can recapitulate the whole epectrum of tumor evolution, from regions with fewer sgRNAs to colonies with substantial sgRNA enrichment (Figures 4E, 4F). Thirdly, Perturb-DBiT co-profiling data enabled identification of perturbation-associated molecular signatures. DEG analysis was conducted between pixels containing sgRNAs for four tumor suppressors validated in this study and pixels containing all other sgRNAs. This revealed significant downregulation (Padj<0.05, Log2FoldChange<-1) of multiple genes known to inhibit tumor growth, along with upregulation (Padj<0.05, Log2FoldChange>1) of a few oncogenes (Figure 4G). This may partially explain why knocking out *SEMA6B*, *MT1E*, *S100A4*, or *ASAH2* may significantly promote tumor colony growth. Further Gene Set Enrichment Analysis (GSEA) (Figure 4H) of the DEGs revealed that knocking out *SEMA6B,* which has been demonstrated as a potential drug target in the prevention and treatment of thyroid carcinoma^59^, appears to facilitate heightened immune response by activating pathways such as T cell activation, regulation of intracellular protein transport, and regulation of cell migration. In summary, Perturb-DBiT enables efficient *in situ* co-profiling of sgRNAs from a genome-wide human CRISPR library alongside whole transcriptomes.

Analysis of the sgRNA data allows for a precise visualization of tumor colonies and rapid identification of functional perturbations on transcriptional state. Concurrently, the transcriptome data analysis reveals genetic modulators that are highly relevant to diverse aspects of tumor biology. For the first time, the integrated analysis of both datasets permits the un-biased spatial uncovering of perturbation-associated molecular signatures using a genome-wide CRISPR library.

### Perturb-DBiT identifies genes modulating the tumor architecture and immune composition of the TME in an immune-competent animal model

We next aimed to use Perturb- DBiT to investigate how different gene perturbations impact TME component. To do this, we transduced E0771 cells, a well-characterized mouse syngeneic cell line, with a pool of lentiviral vectors encoding the genome-wide CRISPR-Cas9 gene knockout library (Brie)^47^.The Brie library comprises a total of 19,674 genes and 78,637 unique sgRNAs. Intravenous injection of E0771-Cas9-GFP-Luciferase Brie library cells into 8-week-old C57BL/6 mice led to tumor colony formation (Figure 5A). After 21 days, lung tissues were collected for Perturb-DBiT PAC co-profiling. Within the ROI (Figure 5B) containing 4 anatomically identifiable major tumors, a total of 91 unique sgRNAs and 17,548 sgRNA UMIs were identified (Table S1, Figure S5A). Gene ontology (GO) analysis of all sgRNA-targeted genes reveals primary involvement in the regulation of cell growth and developmental processes, highlighting significant roles in stem cell proliferation, cell cycle progression, and various embryonic and tissue development stages (Figure S5B). Concurrently, Perturb-DBiT detected an average of 1,995 genes and 4,458 UMIs per pixel (Figure 1B). Cell type-specific marker genes were then identified by unsupervised clustering, uniquely characterizing their expression within each individual cluster for clear distinction from other groups (Figures 5C, S5C). Notably, the distribution of these clusters displayed a strong concordance with tissue morphology (Figure S5D). For example, cluster 4 defined lymph nodes (Figures 5B, 5C), and cluster 0 almost completely overlaps with airway epithelial cells (Figure S5D). Moreover, we observed that two clusters, cluster 1 (C1) and cluster 2 (C2), exhibited conspicuous and distinctive spatial patterns despite their proximity within the same tumor region (Figures 5B, 5C). C1 is defined by DEGs such as *Orc*2, which drives DNA replication^60^ and *St3gal4*, which plays a significant role in tumor progression by promoting neutrophil adhesion to selectins^61^. C2 exhibited upregulation of oncogenes *Pvt*^62^ and *Gse1*^63^, as well as tumor suppressor gene *Cdh13*^64^ (Figures 5D, S5E). However, these distinct genetic profiles alone are not sufficient to definitively differentiate the two tumor clusters.

**Figure 5.**
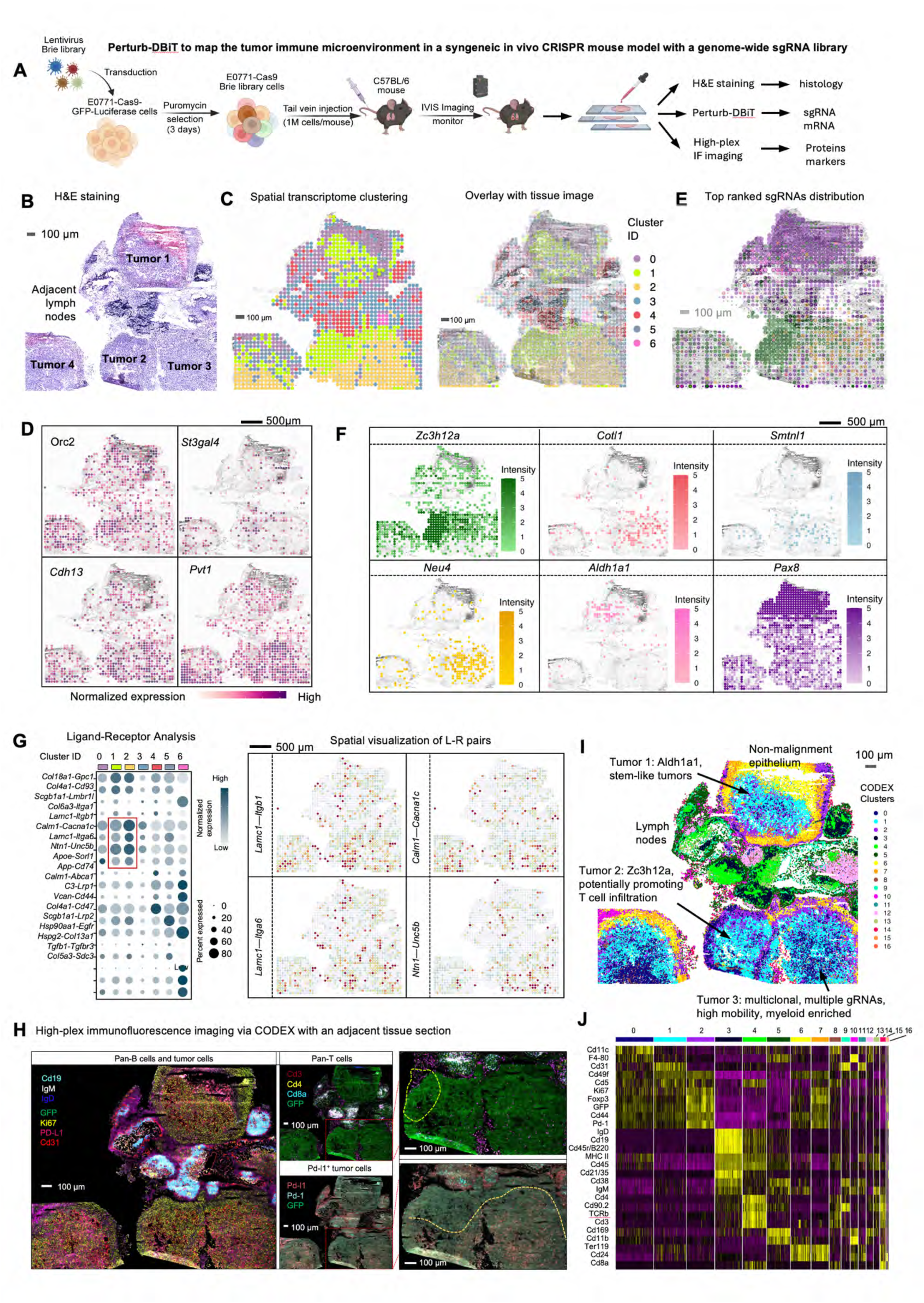
Genome-wide Perturb-DBiT highlights genes that modulate the structural features of tumors and TME. (B) H&E stained image of E0771 syngeneic lung fresh-frozen tissue section with labeling of four anatomically distinct tumor regions and adjacent lymph nodes. (C) Unsupervised clustering of the gene expression matrix revealing 6 distinct clusters overlain on brightfield image of tissue section (right). (D) Spatial intensity maps of four selected genes *Orc2, Cdh13, S13gal4, Pvt1* overlain on brightfield image of tissues section. (E) Spatial intensity plot of top 6 sgRNA hits revealed by Perturb-DBiT overlain on brightfield image of tissue section. (F) Individual spatial intensity plots of top 6 sgRNA hits. **(G)** Left: ligand-receptor interactions within each of the 6 clusters elucidated by unsupervised clustering of gene expression data. Right: spatial maps of four significant ligand-receptor interaction analyses. **(H)** CODEX staining (26 marker panel) is performed on the adjacent tissue section. Left: Pan B cell markers (Cd19, IgM, and IgD), tumor-specific markers (GFP, Ki67, and PD-L1) and vasculature (Cd31). Middle panel, top: background of GFP-positive (tumor) cells and T cells (Cd3+, Cd4+ helper T cells and Cd3+ Cd8a+ cytotoxic T cells). Middle panel, bottom: Tumor cells showing positivity for Pd-l1. Top right: zoomed in image with green circle representing infiltrating region of Cd8+ Tcells and the tumor. Bottom right: zoomed in image with yellow line representing the border of spatial transcriptomics cluster 1 and cluster 2 within the tumor region. (I) UMAP clustering of the CODEX data revealed 17 distinct protein clusters (J) Heatmap showing the top differentially expressed proteins for each cluster

In this instance, we attempt to uncover the mechanisms underlying the distinct characteristics of the two tumor clusters by examining gene perturbations. We selected the top ranked sgRNAs to visualize their spatial distribution (Figures 5E, 5F). Interestingly, we observed a significant enrichment of the sgRNA for *Zc3h12a* within Cluster C1. Previous studies have reported that the deletion of *ZC3H12A* plays a critical role in driving the accumulation of CD8^+^ T cells among tumor-infiltrating lymphocytes^65^, suggesting an enhanced infiltration of CD8+ T cells in the subcluster of C1 proximal to lymph nodes. On the other hand, sgRNA profiles within the C2 exhibit marked heterogeneity. For instance, a majority of sgRNAs that target the *Cotl1* are predominantly found in C2 (Figure 5C). *Cotl1* is known to suppress breast cancer growth by activating the IL-24/PERP pathway and inhibiting non-canonical TGFβ signaling. Consequently, the *Cotl1* perturbation in C2 may result in reduced recruitment of immune cells, while potentially enhancing TGFβ signaling to promote cancer progression^66^.

Although research on Smtnl1’s role in tumors is limited, Keller *et al*.^67^ have demonstrated that SMTNL1 promotes the differentiation of epithelial cells and inhibits their migration in endometrial adenocarcinoma. The presence of enriched sgRNA targeting Smtnl1 in C2 (Figure 5F) could indicate a potential for increased tumor metastasis. NEU4, a sialidase that removes sialic acids from glycoconjugates, has been shown to reduce the motility of hepatocellular carcinoma cells by cleaving sialic acids on CD44, thereby enhancing cell adhesion to the extracellular matrix^68^. Consequently, the preferential enrichment of *Neu4* sgRNA in C2 suggests potential mechanisms contributing to increased cellular motility and elevated levels of tumor metastasis. Furthermore, we observed that two unique sgRNAs, *Aldh1a1* and *Pax8*, are scarce in the C2 but show variable distribution pattern within C1 (Figure 5C, 5F). Specifically, these sgRNAs are rarely found in subcluster of C1 that close to C2, yet they are more prevalent in subcluster of C1 that are distant from C2 (Figure 5C, 5F). Functionally, ALDH1A1 is recognized as a marker for cancer stem cells across various tumor types, suggesting that C2 and adjacent areas of C1 tumor may exhibit enhanced stemness, consistent with the expression levels of the *Pvt1* gene, which is crucial for maintaining robust cancer cell stemness (Figure 5C, 5F)^69^. *Pax8*, typically observed only in high-grade invasive ductal carcinomas, predominantly in triple-negative breast cancer, suggests a comparatively lower invasive potential in the subcluster of C1 tumor that is close to C2 tumor^70^. Overall, C1 tumor appears more heterogeneous, featuring regions of relatively higher CD8^+^ T cell infiltration and variable cancer stem cell-like properties, whereas C2 tumor displays markers of increased aggressiveness and a higher potential for migration or metastasis.

To further examine whether the distinct spatial patterns of sgRNAs observed in C1 and C2 tumors correlate with complex intercellular communication within the TME, we conducted a spatial ligand-receptor (L-R) analysis using the co-profiled transcriptomic data. Differential L-R analysis across all identified spatial populations revealed unique interaction profiles (Figure 5G). Comparing C2 to C1, the top enriched L-R pairs included *Lamc1-Itgb1*, *Calm1-Cacna1c*, *Lamc1-Itga6*, and *Ntn1-Unc5b*. These signaling pathways are known to enhance extracellular matrix (ECM) organization^71^, calcium signaling^72^, tumor cell adhesion to the ECM^73^, and angiogenesis^74^, respectively, which likely contribute to the heightened malignancy observed in C2 tumors. These findings further substantiate the distinct characteristics of C2 tumors as initially revealed by the spatial perturbations analysis via Perturb-DBiT.

Then, to validate the immune infiltration and exclusion in sgRNA-perturbed tumors, we utilized co-detection by indexing (CODEX) with a 27-plex panel, including both immune and tumor cell markers (Table S2), on an adjacent section. Herein, we focus on three anatomically identifiable tumors 1,2, and 3. The CODEX imaging revealed significant overlap with transcriptomic cluster 4 cluster 4 in the lymph nodes, characterized by elevated expression of pan-B cells (Figure 5H). Additionally, the analysis highlighted an increased density of T cells, particularly Cd4^+^ T cells, within the lymph nodes. Despite T cells rarely infiltrating tumor lesions, a notable presence of CD8^+^ T cells was observed near the lymph nodes of the C1 tumor, potentially due to enhanced recruitment facilitated by *Zc3h12a*-specific sgRNA enrichment. In contrast, while PD-L1 expression was nearly undetectable in C1, it was significantly elevated in C2 (Figure 5H), indicating a more pronounced immunosuppressive tumor microenvironment.

CODEX segmentation and clustering has identified distinct cellular clusters, such as pan-B cells (cluster 3), pan-T cells (cluster 4) (Figures 5I, S5F), yet they were limited within the tumor lesions (Figure 5B). Nonetheless, we discerned three major CODEX clusters (Figures 5I, 5J) that co-localize with GFP tumor cells: cluster 0 (CC0), cluster 1 (CC1), and cluster 2 (CC2). Notably, CC1 predominates in C1 tumors, whereas CC0 is scarce; by contrast, C2 features a disproportionately high percentage of CC0. Specifically, compared to CC1, the elevated expression of markers like Cd11c, F4-80, MHC II, Cd45, Cd169, and Cd11b were observed in CC0 (Figure 5I). The markers CD11c and MHC II, associated with myeloid compartment such as dendritic cells (DCs), are essential for antigen presentation and initiating immune responses. However, immunosuppressive mediators secreted by tumor cells could transform potentially immunocompetent DCs into cells that promote immunosuppression, thereby accelerating tumor progression^75^. Similarly, markers such as F4-80 and CD11b indicate macrophages that, within tumors, often differentiate into tumor-associated macrophages (TAMs). TAMs are known to promote tumor growth, angiogenesis, and metastasis, contributing to a more aggressive tumor phenotype^76^. The presence of these cells in high amounts in C2 (Tumor 3) can be linked to a more aggressive tumor phenotype associated with an immunosuppressive myeloid compartment, warranting further validation through more precise marker determination.

Taken together, the results highlight that Perturb-DBiT is highly effective in capturing sgRNAs from immune-competent syngeneic mouse model bearing a whole genome CRISPR library and concurrently sequencing spatial transcriptomics, showcasing its unparalleled capabilities in revealing tumor and tumor microenvironment heterogeneity. Specifically, in the C2 tumor, multiple tumor-related gene perturbations were detected and potentially result in an enrichment of genotypically diverse cell types and cell-cell communication patterns. These are primarily involved in modulating the tumor architecture to promote tumor progression or migration. Moreover, DCs and TAMs may engage in tumor-supporting procedures within the immunosuppressive TME, contributing to the heightened malignancy of the tumors. These dynamics, identified by Perturb-DBiT as pivotal in shaping the tumor architecture and immune composition of the TME, hold potential for developing innovative immunotherapies in the future.

## DISCUSSION

In this study, we introduced Perturb-DBiT, a cutting-edge technology seamlessly integrating spatial transcriptomics with genome-wide spatial CRISPR screening on the same tissue section, offering profound insights into the functional landscape of complex in vivo biological systems. By enabling the simultaneous profiling of genetic perturbations and gene expression across various immune-editing experimental models, Perturb-DBiT enhances our understanding of how genetic modifications influence cellular behaviors within the TME. This versatile platform employs two robust approaches, Perturb-DBiT PAC and DC, to accomplish co-profiling of the biomolecules of interest, accommodating a range of delivery vectors and sgRNA library sizes, and is compatible with both FFPE and fresh frozen samples, making it adaptable for diverse experimental contexts.

This innovative technology enables, for the first time, the unbiased spatial uncovering of perturbation-associated molecular signatures and is compatible with genome-wide CRISPR libraries. By integrating spatial transcriptomics with spatial CRISPR screening, Perturb-DBiT allows researchers to explore the functional landscape of the TME in unprecedented detail. This capability facilitates the identification of specific gene perturbations and their corresponding spatial impacts on cellular neighborhoods. For instance, our findings in the HT29 lung metastatic colonization model reveal distinct clonal architectures, where the spatial colocalization of sgRNAs suggests that they may confer a selective advantage to the tumor cells in which they are expressed, indicating possible synergistic interaction between the genes they target. Further, we have shown that Perturb-DBiT elucidates tumor suppressor and promoter programs, validated by in vitro data. This spatial resolution empowers the discovery of novel molecular pathways and interactions that drive tumor initiation and progression and tumor clonal dynamics, enhancing our understanding of how genetic modifications influence the spatial organization and immune composition within the TME. This comprehensive approach not only broadens the scope of genetic inquiry, but also lays the groundwork for developing targeted therapeutic strategies based on spatial dynamics.

Our analyses have further identified key genes and phenotypic signatures that modulate both tumor architecture and immune composition within the TME. These insights are crucial for understanding the mechanisms of immune evasion and tumor resilience, which are often barriers to effective cancer therapies. By mapping these molecular signatures to specific spatial contexts, we can better understand how functional perturbations in gene expression influence cellular interactions and overall tumor dynamics. The implications of Perturb-DBiT expand beyond cancer research, providing a powerful framework for advancing our understanding of different biological systems. A key advance of Perturb-DBiT that sets this technology apart from traditional single-cell and bulk CRISPR screens as well as current imaging-based spatial screens, lies in its ability to unbiasedly co-profile transcriptome and genome-wide CRISPR screens simultaneously, while maintaining spatial resolution. This comprehensive approach not only broadens the scope of genetic inquiry, but also lays the groundwork for developing targeted therapeutic strategies based on spatial dynamics. By elucidating the functional interplay between genes and their spatial contexts, Perturb-DBiT provides insights that could ultimately lead to improved patient outcome through a more nuanced understanding of the molecular landscape of disease.

## Limitations of the study

A limitation of our current method is yet to realize true single-cell resolution, which may be achieved by fully integrating sequencing-based and imaging-based spatial omics approaches or by further reducing the pixel size^77–80^.To combine the strengths of histological imaging and deep learning, we applied iStar to produce super-resolved single-cell transcriptome data, representing a promising avenue towards this goal. Additionally, the near single-cell resolution can have potential impacts on the spatial capture of sgRNAs including obscurity of spatial heterogeneity in sgRNA distribution impacting exact identification of specific cell populations or subcellular compartments where sgRNA activity is localized. Even with this limitation, this technology provides critical spatial information about sgRNA distribution in an unbiased and base-by-base sequencing fashion which is fully compatible with genome-wide sgRNA libraries.

## RESOURCE AVAILABILITY

### Lead contact

Further information and requests for resources and reagents should be directed to and will be fulfilled by the Lead Contact, Dr. Rong Fan.

### Materials availability

This study did not generate new unique reagents.

### Data and code availability

- Raw and processed sequencing data for this study are in preparation for submitting to the NCBI Gene Expression Omnibus (GEO) database (accession number pending).
- All original code has been deposited at our GitHub space and is publicly available as of the date of publication.
- Any additional information required to reanalyze the data reported in this paper is available from the lead contact upon request.

## ACKNOWLEDGMENTS

We thank Yale West Campus cleanroom team for assistance with microfluidic wafer fabrications and the YPTS team for FFPE tissue sectioning and staining. Computational data analysis was conducted with Yale High Performance Computing clusters (HPC). We acknowledge the support received from the U.S. National Institutes of Health including grants U54CA274509, UH3CA257393, RF1MH128876, U54AG079759, U54AG076043, U01CA294514, R01CA245313, RM1MH132648 (all to R.F.).

## AUTHOR CONTRIBUTIONS

Conceptualization R.F.; data curation A.B., X.T., and P.R. formal analysis, A.B., X.T., P.R., H.L., Z.B., M.Y; writing-original draft, A.B., X.T., Z.B.; writing-review and editing, R.F., S.C.,F.Z.; funding acquisition, R.F.; investigation A.B.,X.T.,P.R.,F.Z., H.L, Z.B, B.T., A.E., H.S., M.Z., W.Z.,T.T.,N.P, M.Y.; methodology A.B, X.T.,P.R.,F.Z.,Z.B., R.F.; resources, F.Z.,P.R., M.Z., H.S.; scientific discussion, Z.B.,B.T.,P.R.,F.Z. and H.L.

## Supplementary Figure Titles and Legends

**Figure S1:** Perturb-DBiT workflow, chemistry design, and optimization, Related to Figure 1

**(A)** Schematic of Perturb-DBiT workflow PAC and DC methods

**(B)** Chemistry design of Perturb-DBiT PAC

**(C)** Chemistry design of Perturb-DBiT DC

**(D)** Optimization of Perturb-DBiT DC primer ratios via bulk experiment on fresh-frozen murine splenic tissue sections.

**Figure S2.** Spatial mapping of sgRNAs in small and medium CRISPR-screening libraries, related to Figure 2

**(A)** Small library guide detection (2 sgRNAs) in Cas9-expressing SMARTA cells in a mouse host spleen using a non-targeting control (NTC) dual-guide. Left, Immunofluorescence imaging of the host tissue is presented with host CD3+ T cells in red, and a blue box marks the ROI used for Perturb-DBiT. Right, The UMI detection of each of the dual guides are shown relative to the ROI. NTC sgRNAs are presented with points (sgRNA1 = red, sgRNA2 = blue) atop a 2D density map (sgRNA1 = blue-to-gold, sgRNA2 = purple-to-yellow).

**(B)** Top: Visualization of top sgRNA hits from 25-micron Perturb-DBiT detection of a medium-sized guide library (288 sgRNAs) in a murine model of autochthonous liver cancer. Bottom: Bar plot of top sgRNA hits from 25-micron Perturb-DBiT applied to a murine model of autochthonous liver cancer

**(C)** Schematics of direct *in vivo* AAV-CRISPR liver screen design.

**(D)** Left: UMAP visualization of pixels based on the dimensional reduction of Mixscape perturbation scores (pUMAP). Non-perturbed (NP) pixels were included in the analysis to serve as a control/reference point. Additionally, pixels were clustered (shown by color and outline) then named by the major perturbations of the cluster (>= 20% of clustered pixels). One cluster with NP and 16 minor perturbations (< 20% of cluster) was named “NP+16”. Right: Violin plot of the expression of 5 selected oncogenes, compared between pUMAP clusters.

**(E)** Volcano plots of the DE analysis between selected pUMAP clusters using pseudo-replicate Wilcox tests. DE genes were those with an adjusted p < 0.05 and an absolute fold-change > 0.1 (up-regulation = red, down- regulation = blue), and the top 8 most significant DE genes were labeled for each analysis.

**(F)** Bubble plot of the top 3 pathways identified from DE genes of each pUMAP cluster (point color). Pathways were detected from gene set enrichment analyses using biological process gene ontologies, and the results are presented by significance (point size) and normalized enrichment score (NES), which provided the enrichment direction and magnitude.

**Figure S3.** Genome-wide high-resolution mapping of HT29 lung metastatic colonization model tissue sections, related to Figure 3

(A) Left: Schematic of Lenti-gRNA Puro construct. Right: Schematic showing the development of HT29 lung metastatic colonization model.

(B) Spatial distribution of selected sgRNAs.

(C) Top ranked DEGs defining each cluster in Figure 3C.

(D) Spatial distribution of identified clusters.

(**E, F**) Super-resolved spatial clustering of the top 3,000 gene profile (E) or all sgRNAs (F) imputed with iStar.

**Figure S4.** Colony formation assays and spatial pseudotime analysis of HT29 lung metastatic colonization model, related to Figure 4

(A) Schematic showing the colony formation assay.

(B) T7E1 assays determined genomic DNA editing and cutting efficiency of each sgRNA.

(C) Colony formation assay analysis utilizing Colony Area plugin in ImageJ of both top enriched sgRNA hits from Perturb-DBiT and validated tumor suppressor/promoter genes for the HT29 lung metastatic colonization model.

(D) Colony formation assay results of validated tumor suppressor/promoter genes for the HT29 lung metastatic colonization model.

**Figure S5.** E0771 syngeneic lung model schematic, sgRNA mapping and GO analysis, and tumor architecture insights, related to Figure 5

**(A)** Schematic of E0771 syngeneic lung model.

**(B)** Spatial intensity plot of sample sgRNA detection via Perturb-DBiT.

**(C)** Top sgRNA hits and their respective counts detected by Perturb-DBiT.

**(D)** GO analysis of top sgRNA hits based on annotation from Ensembl and STRING-db.

**(E)** Heatmap of 6 distinct clusters revealed by unsupervised clustering of spatial transcriptomics data.

**(F)** Top: Pathology annotation differentiating airway epithelial cells and tumor cells within one of the three tumor regions covered by the ROI for Perturb-DBiT.

**(G)** Gene expression spatial plot of *Gse1* overlain on brightfield image of tissue section.

**(H)** CODEX UMAP revealing 16 unique clusters. Cluster 3: B cells, Cluster 4: T cells, Cluster 0,1,2: Tumor cells.

## STAR★METHODS

### KEY RESOURCES TABLE

| REAGENT or RESOURCE | SOURCE | IDENTIFIER |
| --- | --- | --- |
| **Antibodies** |  |  |
| Anti-mouse eno | Bio X Cell | BE0001-1 |
| Human IL-2 | PeproTech | 200-02-50UG |
| Mouse IL-7 | PeproTech | 217-17 |
| Anti-mouse Cd28 (37.51) | Bio X Cell | BE0015-1 |
| Anti-CD3 (2C11) | Bio X Cell | BE0001-1 |
| AF488-labeled B220; clone RA3-6B2 | BioLegend | 103225 |
| AF647-labeled CD3; clone 17A2 | BioLegend | 100209 |
| CODEX antibodies | Akoya Biosciences | See Table S2 |
| CODEX antibodies | Leinco | See Table S2 |
| **Experimental Models** |  |  |
| PD-1 Murine Autochthonous Liver Cancer Model Tissue Sections (CRISPR GEMMs) | Wang, G. et al.<sup>42</sup> | https://doi.org/10.1158/2159-8290.CD-19-1448 |
| LentiX-293T Cell Line | Takara | 632180 |
| E0771 Cell Line | CH3 | 940001 |
| HT29 Cell Line | ATCC | HTB-38 |
| HEK293T Cell Line | ATCC | CRL-1573 |
| C57BL/6J | Jackson Laboratory | 000664 |
| NOD.Cg-Prkdcscid Il2rgtm1Wjl/SzJ (NSG) | Jackson Laboratory | 00557 |
| SMARTA TCR transgenic | Jackson Laboratory | 030450 |
| Rosa26-Cas9 | Jackson Laboratory | 026179 |
| **Bacterial and Virus Strains** |  |  |
| One Shot Stbl3 Chemical Competent E. coli | ThermoFisher | C737303 |
| Endura™ ElectroCompetent Cells | Lucigen | 60242-2 |
| **Chemicals, peptides, and recombinant proteins and DNA** |  |  |
| Acetone | Sigma-Aldrich | 179124-500ML |
| SU-8 photoresist | MicroChem | SU-8 2010 or 2025 |
| Chlorotrimethylsilane | Sigma-Aldrich | 89595-10X1ML |
| Polydimethylsiloxane (PDMS) | Dow Silicones Corporation | GMID:02065622 |
| Tris-HCl pH 8.0 | Sigma-Aldrich | T2694-100ML |
| 5M NaCl | Thermo Fisher Scientific | AM9759 |
| 0.5M EDTA | Life technologies | 15575-038 |
| Xylene | Sigma-Aldrich | 247642-4L-CB |
| Ethanol | Sigma-Aldrich | E7023-500ML |
| Antigen retrieval buffer | abcam | ab93684 |
| Triton X-100 | Sigma-Aldrich | T8787-50ML |
| DPBS | Gibco | 14190-144 |
| Nuclease-free water | Thermo Fisher Scientific | AM9932 |
| RNase Inhibitor | Qiagen | Y9240L |
| E. coli Poly(A) Polymerase | New England Biolabs | M0276L |
| SUPERase•In RNase Inhibitor | Thermo Fisher Scientific | AM2694 |
| Maxima H Minus Reverse Transcriptase | Thermo Fisher Scientific | EP0753 |
| dNTP Mix | Thermo Fisher Scientific | R0192 |
| NEBuffer 3.1 | New England Biolabs | B6003S |
| T4 DNA ligase | New England Biolabs | M0202L |
| SDS | Sigma-Aldrich | 71736-500ML |
| Proteinase K, recombinant, PCR grade | Thermo Fisher Scientific | EO0491 |
| Dynabead MyOne Streptavidin C1 | Thermo Fisher Scientific | 65001 |
| Tris-HCl pH 7.5 | Sigma-Aldrich | T2694-100ML |
| Tween-20 | AmericanBio | AB02038-00500 |
| 20% Ficoll PM-400 | Sigma-Aldrich | F5415-25ML |
| 2X KAPA HiFi HotStart ReadyMix | KAPA Biosystems | KK2602 |
| 20X EvaGreen Plus Dye | VWR International | 76423-470 |
| SPRIselect beads | Beckman Coulter | B23318 |
| Formaldehyde | Thermo Fisher Scientific | BP53125 |
| Lenti-Cas9-GFP | Addgene | 86145 |
| pMD2.G | Addgene | 12259 |
| pCL-Eco | Addgene | 12371 |
| VSV.G | Addgene | 14888 |
| pJR89 | Addgene | 140096 |
| Bpu1102I | Thermo Fisher Scientific | NA |
| BstXI | Thermo Fisher Scientific | NA |
| Bovine serum albumin | Sigma-Aldrich | A7030 |
| DMEM | Gibco | NA |
| Polybrene | Sigma | NA |
| RPMI-1640 | Gibco | NA |
| DAPI Solution | Thermo Fisher Scientific | 62248 |
| T4 DNA Ligase | NEB | M0202L |
| T4 Polynucleotide Kinase | NEB | M0201L |
| **Critical commercial assays** |  |  |
| D5000 ScreenTape Assay | Agilent Technologies | 5067-5588 and 5589 |
| SEQuoia RiboDepletion Kit | BIO-RAD | 17006487 |
| Endotoxin-free maxiprep kits | Qiagen | NA |
| CD4+ T Cell Isolation Kit | Miltenyi Biotec | 130-104-453 |
| **Deposited data** |  |  |
| Raw and processed sequencing data | This paper | GSE link pending |
| Original code | This paper | Github link pending |
| **Oligonucleotides** |  |  |
| Primers, ligation linkers, and DNA barcodes | IDT | See Table S3 |
| **Software and algorithms** |  |  |
| QuPath V0.5.0 | Bankhead, P. et al.<sup>81</sup> | https://qupath.github.io/ |
| Cutadapt V3.4 | Martin, M.<sup>82</sup> | https://cutadapt.readthedocs.io/en/v3.4/ |
| STAR V2.7.7a | Dobin, A. et al.<sup>83</sup> | https://github.com/alexdobin/STAR/releases/tag/2.7.7a |
| ST_Pipeline V1.7.6 | Navarro, J. F. et al.<sup>84</sup> | https://github.com/SpatialTranscriptomicsResearch/st_pipeline |
| GENCODE | Frankish, A. et al.<sup>85</sup> | https://www.gencodegenes.org/ |
| GtRNAdb | Chan, P.P. et al.<sup>86</sup> | https://gtRNAdb.ucsc.edu/ |
| RNAcentral | Petrov, A.I. et al.<sup>87</sup> | https://rnacentral.org/ |
| RStudio V2023.06.2+561 | RStudio | https://posit.co/download/rstudio-desktop/ |
| R V4.3.1 | CRAN | https://www.r-project.org/ |
| R V1.3.1 | CRAN | https://www.r-project.org/ |
| Monocle3 | Cao, J.Y., et al.<sup>58</sup> | https://github.com/cole-trapnell-lab/monocle3 |
| Seurat V4 | Hao, Y. et al.<sup>88</sup> | https://satijalab.org/seurat/ |
| RSeQC package V5.0.1 | Wang, L. et al.<sup>89</sup> | https://rseqc.sourceforge.net/ |
| miRBase | Kozomara, A. et al.<sup>90</sup> | https://www.mirbase.org/ |
| Integrative Genomics Viewer (IGV) | Robinson, J.T. et al.<sup>91</sup> | https://igv.org/ |
| rMATs-turbo pipeline V4.1.2 | Shen, S. et al.<sup>92</sup> | https://github.com/Xinglab/rmats-turbo |
| REDportal V2.0 | Picardi, E. et al.<sup>93</sup> | http://srv00.recas.ba.infn.it/atlas/ |
| samtools V1.16.1 | Danecek, P. et al.<sup>94</sup> | https://github.com/samtools/samtools |
| Bowtie(v.1.3.0) | Langmead, et al.<sup>95</sup> | https://github.com/BenLangmead/bowtie |
| iStar | Zhang D. et al.<sup>96</sup> | https://github.com/daviddaiweizhang/istar |
| bedtools V 2.31.0 | Quinlan, A. R. et al.<sup>97</sup> | https://github.com/arq5x/bedtools2 |
| scVelo | Bergen, V. et al.<sup>98</sup> | https://scvelo.readthedocs.io/en/stable/ |
| Strelka V2.9.10 | Kim, S. et al.<sup>99</sup> | https://github.com/Illumina/strelka |
| BWA V0.7.17 | Li, H. et al.<sup>100</sup> | https://github.com/lh3/bwa |
| GATK V4.2.0.0 | Data Sciences Platform at the Broad Institute | https://gatk.broadinstitute.org/hc/en-us/sections/4405443482011-4-2-2-0 |
| ggtree V3.8.2 | Xu, S. et al.<sup>101</sup> | https://www.bioconductor.org/packages/release/bioc/html/ggtree.html |
| inferCNV V1.16.0 | inferCNV of the Trinity CTAT Project | https://github.com/broadinstitute/inferCNV/wiki |
| CNVkit V0.9.11 | Talevich, E. et al.<sup>102</sup> | https://cnvkit.readthedocs.io/en/stable/ |
| Connectome V1.0.0 | Raredon, M. S. B. et al.<sup>103</sup> | https://msraredon.github.io/Connectome/ |
| FGSEA | Korotkevich, G. et al.<sup>104</sup> | https://github.com/ctlab/fgsea/ |
| Prism V9 | GraphPad | https://www.graphpad.com/ |
| Bwa mem v.0.7.17 | Li, H. et al.<sup>100</sup> | https://github.com/lh3/bwa |
| VarScan v2.3.9 | Koboldt, D.C., et al.<sup>105</sup> | [https://github.com/dkoboldt/varsan/releases](https://github.com/dkoboldt/varscan/releases) |
| **Other** |  |  |
| Poly-L-Lysine Slide | Electron Microscopy Sciences | 63478-AS |
| Chrome photomask | Front Range Photomasks | N/A |
| Silicon wafer | WaferPro | C04004 |
| EVOS M7000 Imaging System | Thermo Fisher Scientific | AMF7000 |
| NIS-Elements software | Nikon Instruments | Ver.5.42.04 |
| DNA low-bind tube | Eppendorf SE | 022431021 |
| TapeStation 4150 | Agilent Technologies | G2992AA |
| NovaSeq 6000 Sequencing System | Illumina | N/A |
| CODEX PhenoCycler-Fusion | Akoya Biosciences | N/A |
| Leica TCS SP5 Confocal microscope | Leica Microsystems | N/A |
| CFX Connect Real-Time System | BIO-RAD | N/A |

### EXPERIMENTAL MODEL AND SUBJECT DETAILS

#### Mice: SMARTA TCR transgenic and B16-OVA tumor models

Mice were housed and bred at the St. Jude Children’s Research Hospital Animal Resource Center in specific pathogen-free conditions. C57BL/6, *Rosa26-Cas9*^41^ and SMARTA TCR transgenic^106^ mice were purchased from the Jackson Laboratory. Female and male mice were used at 6–10 weeks of age. Mice were housed and bred at the St. Jude Children’s Research Hospital Animal Resource Center in specific pathogen-free conditions. The research conducted in this study complied with all the relevant ethical regulations. Experiments and procedures were approved by and performed in accordance with the Institutional Animal Care and Use Committee (IACUC) of St. Jude Children’s Research Hospital. *Rosa26-Cas9* transgenic mice were injected subcutaneously with 1 × 10^6^ B16-OVA-Cas9 cells expressing the CRISPR sgRNA library in the right flank. Tumor tissues were harvested 14 days after implantation. Tumor size limits were approved to reach a maximum of 3,000 mm^3^ or ≤ 20% of body weight (whichever was lower) by the IACUC of St. Jude Children’s Research Hospital.

#### Mice: CRISPR GEMM, E0771 syngeneic lung colonization, and HT29 lung metastatic colonization models

For CRISPR GEMM model of liver cancer: mixed gender (randomized males and females) 8–12 week old Rosa26-LSL-Cas9–2A-EGFP (LSL-Cas9) mice were bred with C57BL/6J mice, FVB.129S6(B6)- Gt(ROSA)26Sortm1(Luc)Kael/J mice, or C57BL/6N-Gt(ROSA)26Sortm13(CAG-MYC,-CD2*)Rsky/J mice^42^. For HT29 lung metastatic colonization model: 8-week-old NSG mice were used. For E0771 syngeneic lung tumors: 8-week-old female C57BL/6J mice were used. This study operated under institutional regulatory approval. All work was performed under the guidelines of Yale Environment, Health and Safety (EHS) Committee with an approved protocol. Animal work for CRISPR GEMM model of liver cancer, HT29 lung metastatic colonization model and E0771 syngeneic lung tumors operated under the guidelines of Yale University Institutional Animal Care and Use Committee (IACUC) with approved protocols. Please refer to previously published work for method details^42^.

#### Mouse paraffin tissues

Fresh tissue sections were obtained from the mentioned mouse specimens. These sections were preserved in PFA and subsequently prepared for paraffin embedding. To maintain RNA, DNA, and protein integrity, all procedures from tissue collection to embedding were carried out in a contamination-free environment. The stained tissue sections were then examined by experienced histologists to confirm their suitability for further analysis.

#### Sample handling and tissue section preparation

Paraffin blocks from and mouse samples were sliced into 7-10 micrometer-thick sections. These sections were then placed on glass slides coated with Poly-L-Lysine. Additional serial sections were collected simultaneously for Perturb-DBiT. While the sectioning of mouse formalin-fixed paraffin-embedded (FFPE) samples was performed at the YPTS facility, the sectioning of mouse frozen (FF) samples was carried out in-house. The prepared paraffin sections were subsequently stored at a temperature of -80 degrees Celsius until required for further analysis.

### METHOD DETAILS

#### B16-OVA tumor model: cell lines and retroviral library transduction

B16-OVA cell line was provided by D. Vignali (University of Pittsburgh). The HEK293T cell line was purchased from the American Type Culture Collection (ATCC). The Plat-E cell line was provided by Y.-C. Liu (La Jolla Institute of Immunology). All cell lines were cultured in Dulbecco’s modified essential medium (DMEM) (Gibco) or RPMI-1640 medium (Gibco) supplemented with 10% (*v*/*v*) FBS and 1% (*v*/*v*) penicillin-streptomycin at 37°C with 5% CO2. No commonly misidentified cell lines (International Cell Line Authentication Committee) were used in this study. Cell lines were tested and determined to be free of mycoplasma contamination. The aforementioned cell lines were not independently authenticated. To generate Cas9-expressing tumor cell line B16-OVA-Cas9, lentivirus was produced by co-transfecting Lenti-Cas9-GFP (86145, Addgene) plasmid with psPAX2 (12260, Addgene) and pMD2.G (12259, Addgene) packing plasmids into HEK293T cells. The supernatant containing viral particles was harvested 48 hours after transfection. B16-OVA cells were transduced with viral supernatant for 48 hours in RPMI-1640 (for B16-OVA) + 10% (*v*/*v*) FBS supplemented with 10 μg/ml polybrene (Sigma), followed by sorting of transduced (GFP^+^) into single clones, followed by expansion. Cas9 expression was verified by immunoblot analysis^107^. To generate tumor cells transduced with the retroviral CRISPR sgRNA library (LMA-DC-EFS vectors used as backbone) targeting epigenetic genes, retrovirus was produced by co-transfecting the retroviral library with pCL-Eco (12371, Addgene) and VSV.G (14888, Addgene) packing plasmids into Plat-E cells. The supernatant containing viral particles was harvested 48 hours after transfection. B16-OVA-Cas9 cells were transduced with viral supernatant for 48 hours in RPMI 1640 (for B16-OVA) + 10% (*v*/*v*) FBS supplemented with 10 μg/ml polybrene (Sigma), followed by sorting Ametrine^+^ cells. Cells were cultured for another 14 days for genome editing and expansion.

#### B16-OVA tumor model: Dual-guide direct-capture retroviral library construction

For the curated gene list containing 38 epigenetic genes, a total of two gRNA sequences distributed on an individual construct were designed for each gene. To construct the library, a customized oligonucleotide pool containing 76 oligonucleotides targeting those 38 epigenetic genes and 10 NTCs (each oligonucleotide contains two guides targeting the same gene or NTC) (Table S4) was ordered from Twist Biosciences. The oligonucleotide design follows the overall structure: 5′-PCR adapter-CCACCTTGTTGG-protospacer A– GTTTCAGAGCAGTCTTCGTTTTCGGGGAAGACAAGAAACATGG-protospacer B–GTTTAAGAGCTAAGC-PCR adapter-3′. The dual-guide library was generated using a two-step cloning strategy as previously described^108^. In brief, the PCR-amplified oligonucleotide pool was digested with BstXI and Bpu1102I (Thermo Fisher) and ligated into a similarly digested LMA-DC-EFS vector. The ligation product was then electroporated into Endura Duos (Lucigen) and amplified, and the resulting intermediate library was assessed for quality using next generation sequencing (NGS). For quality control, sgRNA skewing was measured using the script calc_auc_v1.1.py^109^ to monitor how closely sgRNAs are represented in a library, and sgRNA distribution was plotted with the area under the curve < 0.7 to pass quality control. The Python script count_spacers.py^110^ was used as an additional measure for quality control. Next, the CR3^cs1^-hU6 insert from pJR89 (140096, Addgene) was isolated by digestion with BsmBI followed by gel extraction. The intermediate library from above was digested with BbsI and treated with rSAP. Finally, the CR3^cs1^-hU6 insert was ligated into the intermediate library vector, purified by isopropanol purification and electroporated into Endura Duos. Electroporated cells were plated overnight at 32 °C, collected the next day and the plasmid library extracted using endotoxin-free maxiprep kits (Qiagen). The amplified library was then validated by NGS as described above.

#### SMARTA TCR transgenic model: sgRNA transduction in CD4^+^ T cells and adoptive transfer after LCMV infection

Naive Cas9-expressing SMARTA cells were isolated from the spleen and peripheral lymph nodes of Cas9- SMARTA mice using the naive CD4^+^ T cell isolation kit according to the manufacturer’s instructions. Purified naive SMARTA cells were activated *in vitro* for 18 h with 5 μg/ml anti-CD3 and 5 μg/ml anti-CD28 before viral transduction. Viral transduction was performed by spin-infection at 800*g* at 25 °C for 3 h with 10 μg/ml polybrene (Sigma). Cells were then cultured with human IL-2 or mouse IL-7 for 4 days. Transduced cells were sorted based on the expression of Ametrine using a Reflection (i-Cyt) and then adoptively transferred into C57BL/6 recipients 1 day after infection of the host mice with LCMV. For LCMV infection, 2 × 10^5^ plaque- forming units (PFU) of the Armstrong strain of LCMV were injected intraperitoneally. sgRNAs were designed using an online tool (https://portals.broadinstitute.org/gpp/public/analysis-tools/sgrna-design). sgRNAs used in this study were cloned into LMA-DC-EFS dual-guide direct-capture retroviral sgRNA vectors described before^37^. The spacer sequences for the two non-targeting control sgRNAs on the dual-guide retroviral vector: 5’- AGGACTATCCGCGGGATTAG -3’ and 5’- ATGACACTTACGGTACTCGT -3’; At day 6 post LCMV infection, spleens were harvested for section.

#### Lentivirus production for HT29 lung metastatic colonization model and E0771 syngeneic model

For lentivirus production, 20 μg of lentiviral vectors—such as Lenti-EF1a-GFP-T2A-Luciferase-WPRE, Lenti- EF1a-Cas9-Blasticidin-WPRE, and either a target sgRNA or library in Lenti-U6-sgRNA-EFS-Puro-WPRE, were co-transfected with 10 μg of pMD2.G and 15 μg of psPAX2 into LentiX293T cells. This co-transfection was carried out in a 150 mm dish at 80-90% confluency using 135 µL of lipoD293 transfection reagent. Virus-containing supernatant was harvested at 48 hours post-transfection, then centrifuged at 1,500 g for 15min to remove cellular debris. The virus was subsequently concentrated using a LentiX concentrator, aliquoted, and stored at -80°C. The library virus was titrated by infecting HT29 or E0771 cells and applying a selection pressure of 3 μg/ml puromycin.

#### Generation of human Brunello library or mouse Brie library in transduced cells

HT29 or E0771 cells stably expressing GFP and luciferase were created by transducing them with lentivirus EF1a-GFP-T2A-Luciferase-WPRE. Cells expressing GFP were isolated using flow cytometry sorting.

Subsequently, HT29-GFP-Luciferase or E0771-GFP-Luciferase cells stably expressing Cas9 were produced by transduction with lentivirus EF1a-Cas9-Blasticidin-WPRE, followed by a selection period of 7 days using 20 µg/ml blasticidin. The Brunello library (Table S4) was transduced into HT29-Cas9 cells, while the Brie library (Table S4) was used to infect E0771-Cas9 cells. The sgRNA libraries were transduced at a calculated MOI of 0.3, with a minimal representation of over 200x transduced cells per sgRNA. Post-infection, cells were incubated at 37°C for 24 hours before selection in media containing 3 µg/ml puromycin for 3 days. The successfully transduced cells were then injected into mice.

#### HT29 lung metastatic colonization model

1 x 10^^6^ HT29-Cas9-Brunello library cells were intravenously injected into 8-week-old NSG mice. The formation of metastases was monitored using IVIS imaging.

#### E0771 syngeneic lung colonization model

1 x 10^^6^ E0771-Cas9-Brie library cells were intravenously injected into 8-week-old female C57BL/6J mice. The formation of metastases was also monitored using IVIS imaging.

#### Clonogenic assay of HT29 cells in vitro

HT29-Cas9 cells were transduced with either a non-targeting control (NTC), AAVS1 sgRNA control, or target gRNA lentivirus (Lenti-EF1a-gRNA-EF1a-Puro). The gRNA-transduced cells underwent selection via puromycin for 3 days, followed by validation of sgRNA genome editing using the T7E1 assay. For the tumor cell clonogenic assay, cells were dissociated into a single-cell suspension. This suspension was diluted and seeded at densities of 5,000 cells per well in a 6-well plate. After 14 days, the medium was removed, and the cells were gently rinsed with PBS, fixed with 75% ethanol, and stained with 0.5% crystal violet. Cell colonies were analyzed using the Colony Area plugin in ImageJ, which automatically quantifies colony formation in clonogenic assays^111^. Gene oligos used for assay are listed in table (Table S5)

#### Microfluidic Device Fabrication for Perturb-DBiT

The microfluidic device was fabricated using a standard soft lithography process, detailed in our previous publication^34^. In summary, customized high-resolution photomasks were printed and procured from Front Range Photomasks (Lake Havasu City, AZ). Photomasks were then cleaned with acetone to remove contaminants and master wafers were created using SU-8 negative photoresist (SU-2010 or SU-2025) on silicon wafers. Fabrication widths varied between 10, 20, and 50 microns. The wafers were treated for 20 minutes with chlorotrimethylsilane to achieve high-fidelity hydrophobic surface. Polydimethylsiloxane (PDMS) microfluidic chips were produced using a 10:1 mixture of base and curing reagents (following the manufacturer’s protocol) and poured over the master wafers. The mixture was degassed in a vacuum for 30 minutes and subsequently cured at 70°C for a minimum of 2 hours. Upon solidification, the PDMS devices were cut out and inlets and outlets were punched prior to Perturb-DBiT application.

#### DNA barcodes annealing

The DNA oligos used in this study were sourced from Integrated DNA Technologies (IDT, Coralville, IA) with the sequences provided in Table S3. Barcode (100 µM) and ligation linker (100 µM) were mixed in equal parts (1:1) in a 2X annealing buffer (20 mM Tris-HCl pH 8.0, 100 mM NaCl, 2 mM EDTA). The annealing process followed this thermal cycling protocol: 95°C for 5 min, gradual cooling to 20°C at a rate of −0.1°C/s, and then a 3-minute hold at 12°C. The annealed barcodes can be preserved at −20°C until use.

#### FFPE tissue deparaffinization and decrosslinking

A tissue section was retrieved from −80°C storage and allowed to sit at room temperature for 10 minutes until moisture had evaporated. The tissue slide was then baked at 1-2 hours at 60°C to soften and melt the paraffin. Paraffin was removed by immersing the slides into two changes of xylene, followed by rehydration through a graded series of ethanol solutions for five minutes each, including two rounds of 100% ethanol and one round each of 90%,70% and 50% ethanol, with a final rinse of distilled water. The slide was then completely covered by a pre-boiled 1X antigen retrieval buffer (on a hot plate) for 10 minutes followed by a 15-minute cooldown to room-temperature on ice. After a brief rinse in distilled water, the tissue was scanned using a 10x objective on the EVOS M7000 Imaging System to capture the intact tissue. Sections could either be used right away for Perturb-DBiT, or could be stored for up to 4-weeks at −80°C.

#### Perturb-DBiT DC permeabilization, reverse transcription, and spatial barcoding with microfluidic devices

The tissue was permeabilized at room temperature for 20 min with 1% Triton X-100 in DPBS, followed by a 0.5X DPBS-RI (1X DPBS diluted with nuclease-free water, 0.05 U/µL RNase Inhibitor) wash to stop permeabilization. The tissue slide was then air-dried, imaged, and equipped with a PDMS reservoir for bulk reverse transcription. 62.2uL of the reverse transcription mix, using a 1:1 ratio of sgRNA: mRNA capture reagents (10 µL 25 µM Transcriptome RT Primer, 10 µL 25 µM CRISPR RT Primer, 20 µL 0.5X DPBS-RI, 12 µL 5X RT Buffer, 6 µL 200U/µL Maxima H Minus Reverse Transcriptase, 3 µL 10mM dNTPs, 0.8 µL 20 U/µL SUPERase•In RNase Inhibitor, 0.4 µL 40 U/µL RNase Inhibitor) was added into the PDMS reservoir and sealed with parafilm prior to incubation in a wet box for 30 min at room temperature. The sample was then incubated for 90 min at 42°C, followed by a 5 min 50mL DPBS shake-wash.

To ligate barcode A and C *in situ*, the first PDMS device was positioned on top of the tissue slide, ensuring alignment of the center channels with the ROI. The device was then imaged to record its positioning on the tissue for downstream analysis and alignment. Next, an acrylic clamp was applied to firmly secure the PDMS device to the tissue slide, preventing reagents from leaking between the microfluidic channels. The ligation mix (100 µL 1X NEBuffer 3.1, 11.3 µL nuclease-free water, 26 µL 10X T4 ligase buffer, 15 µL T4 DNA ligase, 5 µL 5% Triton X-100, 2 µL 40 U/µL RNase Inhibitor, and 0.7 µL 20 U/µL SUPERase•In RNase Inhibitor) was then prepared on ice. For the barcoding reaction, a total of 5 µL ligation solution (3 µL ligation mix and 1 µL each of barcode A and C) was carefully pipetted into each of the 50 inlets of the device at room temperature. The ligation solution within each of the 50 inlets was then gently pulled throughout each of the 50 channels of the device using a delicately adjusted vacuum. After incubation in a wet box for 30 min at 37°C, the first PDMS device was removed, and the slide was washed with 50mL DPBS. The second PDMS device, featuring 50 channels perpendicular in direction to the first PDMS device, was then attached to the tissue slide with careful alignment of the central channels as-to finalize the positioning of the ROI. Following another microscopic image for downstream alignment, the acrylic clamp was applied once again to firmly secure the PDMS device in place and the ligation of barcode C and D was similarly performed. Following barcoding, five flow-washes with 500 µL nuclease-free water were performed and the final scan was conducted to record the tissue-imprinted microchannel marks, making clear the ROI.

### Perturb-DBiT PAC permeabilization, *in situ* polyadenylation, reverse transcription, and spatial barcoding with microfluidic devices

The tissue was permeabilized and washed as described above. The tissue was then air-dried and equipped with a PDMS reservoir surrounding the ROI. *In Situ* polyadenylation was performed using *E.Coli* Poly(A) Polymerase. Samples were first equilibrated incubating 100 µL poly(A) wash buffer (88 µL nuclease-free water, 10 µL 10X Poly(A) Reaction Buffer, 2 µL 40 U/µL RNase Inhibitor) at room temperature for 5 min. After removing the Poly(A) wash buffer, 60 µL of the Poly(A) enzymatic mix (38.4 µL nuclease-free water, 6 µL 10X Poly(A) Reaction Buffer, 6 µL 5U/µL Poly(A) Polymerase, 6 µL 10mM ATP, 2.4 µL 20 U/µL SUPERase•In RNase Inhibitor, 1.2 µL 40 U/µL RNase Inhibitor) was added to the reservoir and incubated in a wet box at 37°C for 25 minutes. Tissue slides were then shake-washed in DPBS as described above before adding 60.2uL of the reverse transcription mix(10 µL 25 µM Transcriptome RT Primer, 20 µL 0.5X DPBS-RI, 12 µL 5X RT Buffer, 6 µL 200U/µL Maxima H Minus Reverse Transcriptase, 8 µL nuclease-free water, 3 µL 10mM dNTPs, 0.8 µL 20 U/µL SUPERase•In RNase Inhibitor, 0.4 µL 40 U/µL RNase Inhibitor) to the PDMS reservoir with a similar incubation as described above. To ligate barcode A *in situ*, the first PDMS device was positioned and imaged as previously described. The ligation mix (100 µL 1X NEBuffer 3.1, 61.3 µL nuclease-free water, 26 µL 10X T4 ligase buffer, 15 µL T4 DNA ligase, 5 µL 5% Triton X-100, 2 µL 40 U/µL RNase Inhibitor, and 0.7 µL 20 U/µL SUPERase•In RNase Inhibitor) was then prepared on ice. For the barcoding reaction, a total of 5 µL ligation solution (4 µL ligation mix and 1 µL of barcode A) was carefully pipetted into each of the 50 or 100 inlets of the device at room temperature. The ligation solution within each of the 50 or 100 inlets was then gently pulled throughout each of the 50 or 100 channels of the device using a delicately adjusted vacuum. After incubation in a wet box for 35 min at 37°C, the first PDMS device was removed, and the slide was washed as described above. The second PDMS device, featuring 50 or 100 channels perpendicular in direction to the first PDMS device, was then similarly attached and imaged. The ligation of barcode B was similarly performed.

Following barcoding, five flow-washes with 500 µL nuclease-free water were performed and the final scan was conducted to record the tissue-imprinted microchannel marks, making clear the ROI.

#### Tissue lysis and extraction of cDNA

For both Perturb-DBiT DC and PAC, the ROI of the barcoded tissue was surrounded with a clean PDMS reservoir and clamped securely with an acrylic chip. A 2X lysis buffer (20 mM Tris-HCl pH 8.0, 400 mM NaCl, 100 mM EDTA, and 4.4% SDS) was prepared ahead of time. 110 µL of lysis mix (50 µL 1X DPBS, 50 µL 2X lysis buffer, 10 µL 20 µg/µL Proteinase K solution) was then added to the reservoir and sealed with parafilm prior to a 2 hour incubation at 55°C in a wet box. Following the reaction, 110 µL sgRNA and mRNA cDNA-rich liquid was collected into a 1.5mL DNA low-bind \The tissue lysate could then be stored at -80°C prior to the next steps.

#### Purification of cDNA and template switching

The 100 µL sgRNA and mRNA cDNA was purified using 40 µL of Dynabeads MyOne Streptavidin C1 beads resuspended in 100 µL of 2X B&W buffer (10 mM Tris-HCl pH 7.5, 1 mM EDTA, 2 M NaCl) and incubated with rotation at room temperature for 60 min to ensure sufficient binding. Following magnetic separation and two washes with 1X B&W buffer (with 0.05% Tween-20) and an additional two washes with 10 mM Tris-HCl pH 7.5 containing 0.1% Tween-20, cDNA molecules bound with streptavidin beads were then resuspended in 200 µL of TSO Mix (75 µL nuclease-free water, 40 µL 5X RT buffer, 40 µL 20% Ficoll PM-400, 20 µL 10mM dNTPs, 10 µL 200U/µL Maxima H Minus Reverse Transcriptase, 5 µL 40 U/µL RNase Inhibitor, 10 µL 100 µM TSO Primer). Template switching was performed with rotation at room temperature for 30 min and then at 42°C for another 90 min. Beads then went through two washes, one with 10mM Tris-HCl pH 7.5 containing 0.1% Tween-20 and another with nuclease-free water. The beads were then PCR amplified according to different protocols, depending on if Perturb-DBiT DC or PAC was employed.

#### Perturb-DBiT DC PCR amplification and library preparation

Washed beads were then resuspended in 200 µL of PCR mix with a 1:1 ratio of sgRNA: mRNA capture primers (100 µL 2X KAPA HiFi HotStart ReadyMix, 76 µL nuclease-free water, 8 µL 10 µM PCR Primer 1, 8 µL 10 µM PCR Primer 2, 8 µL 10 µM PCR Primer 3). This mixture was distributed into 4 PCR strip tubes and the following PCR program was used: 95°C for 3 min, with five cycles at 98°C for 20 s, 63°C for 45 s, 72°C for 3 min, followed by an extension at 72°C for 3 min and 4°C hold. After magnetically removing beads, 19 µL of the PCR solution was combined with 1 µL 20X EvaGreen for quantitative real-tiyme PCR (qPCR) using the same program while the remaining solution underwent normal PCR simultaneously. The cycle numbers were finally determined when ½ of the saturated signal was observed. The full PCR product was then purified using a 0.6X ratio of SPRIselect beads while saving the supernatant. The pellet portion (transcriptomic cDNA) adhered to the standard manufacturer’s instructions for purification while 150 µL supernatant (sgRNA cDNA) underwent a second 1.2X SPRI selection. Following a second elution in 50 µL nuclease-free water, a 1.0X Spri selection was performed with the transferred supernatant. Finally, sgRNA product was eluted in 30 µL nuclease-free water. The transcriptomic cDNA and sgRNA each underwent analysis using a TapeStation system with D5000 DNA reagents and ScreenTape. Depending on experiment-specific sgRNA cDNA QC, sgRNA enrichment was necessary. We prepared a sgRNA enrichment PCR mix (50 µL 2X KAPA HiFi HotStart ReadyMix, 2ng sgRNA product from the previous reaction, 8 µL 10 µM PCR Primer 2, 8 µL 10 µM CRISPR PCR Primer 3, 5uL 20X EvaGreen nuclease-free water to 100uL) using a 100uL system and performed qPCR using the same program. After enrichment, sgRNA underwent a 1.2X SPRI cleanup prior to elution in 30 uL nuclease-free water. After ensuring high-quality cDNA, we performed direct ligation library preparation for sequencing. We prepared two solutions, each consisting of 50 µL 2X KAPA HiFi HotStart ReadyMix, ∼2ng sgRNA or mRNA cDNA, 4 µL 10 µM P5 Primer, and 4 µL 10 µM P7 Primer or CRISPR P7 Primer for transcriptome or sgRNA library, respectively. We then performed qPCR for each sample using the aforementioned program to build libraries. The transcriptome library then underwent purification using a 0.8X SPRI ratio, while the sgRNA library then underwent purification using a 1.2X SPRI ratio. Both libraries were quality control checked using TapeStation and then were sequenced on an Illumina NovaSeq 6000 Sequencing System with a paired-end 150bp read length.

#### Perturb-DBiT PAC PCR amplification, rRNA removal, and library preparation

For the Perturb-DBiT PAC method, washed beads were then resuspended in 200 µL of PCR mix (100 µL 2X KAPA HiFi HotStart ReadyMix, 84 µL nuclease-free water, 8 µL 10 µM PCR Primer 1, and 8 µL 10 µM PCR Primer 2). This mixture was distributed into 4 PCR strip tubes and the following PCR program was used: 95°C for 3 min, with 13 cycles at 98°C for 20 s, 63°C for 45 s, 72°C for 3 min, followed by an extension at 72°C for 3 min and 4°C hold. The full PCR product was then purified using a 0.7X ratio of SPRIselect beads while saving the supernatant. The pellet portion (transcriptomic cDNA) adhered to the standard manufacturer’s instructions for purification and was eluted in 20 µL nuclease-free water while the transferred supernatant (containing sgRNA) underwent a second 1.2X SPRI selection before being eluted in 20 µL nuclease-free water.

The resulting mRNA cDNA and sgRNA underwent analysis using a TapeStation system with D5000 DNA reagents and ScreenTape. The SEQuoia Ribodepletion Kit was then used to eliminate rRNA-derived fragments and mitochondrial rRNA from the amplified sgRNA and cDNA product, following manufacturer’s guidelines. Based on the readout from TapeStation analysis, 20ng of sgRNA and mRNA cDNA was used as an input amount and two rounds of rRNA depletion were performed. Next, the aforementioned PCR program was executed on the two separate samples for 7 cycles to directly ligate sequencing primers using a 100 µL system (50 µL 2X KAPA HiFi HotStart ReadyMix, ∼42 µL solution from the rRNA removal step, 4 µL 10 µM P5 Primer, and 4 µL 10 µM P7 Primer). The final transcriptomic library underwent 0.7X SPRI selection, while the final sgRNA library underwent 0.6X SPRI selection followed by a 1.2X SPRI selection. The two libraries were checked for quality control using TapeStation and then were sequenced on an Illumina NovaSeq 6000 Sequencing System with a paired-end 150bp read length.

#### CODEX spatial phenotyping using PhenoCycler-Fusion

The CODEX PhenoCycler-Fusion protocol ([Link to protocol](https://www.akoyabio.com/wp-content/uploads/2021/01/CODEX-User-Manual.pdf)) for fresh frozen tissue sections from Akoya Biosciences was followed. The tissue slide was removed from the -80C freezer and placed on drierite beads for 5 minutes. After this, the tissue slide was immersed in acetone for 10 mins and then allowed to dry for 2 minutes. The tissue slide was then passed through a series of hydration steps – immersed in hydration buffer for 2 minutes twice, followed by fixation for 10 minutes using 1.6% PFA. Finally, the tissue section was immersed in staining buffer for 30 minutes and the antibody cocktail prepared. The tissue slide was incubated with the antibody cocktail at room temperature for 3 hours in a humidity chamber. After incubation, the tissue underwent a series of steps including post-fixation, ice-cold methanol incubation, and a final fixation step.

Attached to the flow cell, the tissue section was incubated in 1X PhenoCycler buffer with additive for at least 10 minutes to improve adhesion. The CODEX cycles were then set up, the reporter plate was prepared and loaded, and the imaging process began. A final qptiff file was generated, at the end which could be viewed using QuPath V0.5^81^. Specific details regarding the Phenocycler reagents, volumes, panels, and cycles can be found in Table S2.

#### Immunofluorescent Staining

Immunofluorescence analysis of spleen tissue sections was performed following rehydration in PBS, and subsequent blocking of non-specific binding using blocking buffer comprised of PBS containing 1% BSA, 0.05% Tween-20 and 5% normal donkey serum. Slides were incubated overnight at 4°C in PBS containing 1% BSA and AF488-labeled B220, AF647-labeled CD3. Slides were washed in PBS and mounted in Vectashield vibrance mounting media containing DAPI (Vector labs, H-1800). Images were acquired an A1RHD resonant scanning confocal comprised of a Ti2-E inverted microscope equipped with an A1RHD25 scanner detector, Plan Apo 20X 1.0 NA objective, laser lines as appropriate for 405/488/647 illumination and detection and were analyzed using NIS-Elements software (Nikon Instruments; version 5.42.04).

### QUANTIFICATION AND STATISTICAL ANALYSIS

#### Sequence alignment and mRNA expression matrix generation

The FASTQ file Read 2 underwent processing from the raw sequencing data, involving UMI and spatial barcode extraction. Read 1 containing cDNA sequences was trimmed using Cutadapt V3.4^82^ and then aligned to either the mouse GRCm38-mm10 or human GRCh38 reference genome using STAR V2.7.7a^83^. Utilizing ST_Pipeline V1.7.6^84^, spatial barcode sequences were demultiplexed based on the predefined microfluidic coordinates and ENSEMBL IDs were converted to gene names. This in turn generated the gene-by-pixel expression matrix for downstream analysis. Matrix entries corresponding to pixel positions with no underlying tissue were excluded for analysis and missing pixels are inferred from nearby data to facilitate clustering analysis across the entire mapped area.

### Gene expression analysis

Gene expression (GEX) data were loaded into a Seurat object along with spatial positions. The GEX data were normalized and scaled using SCTransform method with the “v2” strategy for variance-stabilizing transformation^112^. The data were then dimensionally reduced by principal components analysis (PCA), followed by UMAP (min.dist = 0.01) using the PCA components that explained the majority of the data variance (chosen by elbow plot method) ^14,113^. Pixels were clustered by constructing a shared nearest-neighbors (SNN) graph from PCA data and using the FindClusters function (algorithm = 3). The optimal clustering resolution for modularity optimization was chosen based on the minimal total within-cluster sum-of-squares and corresponding maximal average silhouette-width when considering > 3 clusters.

Differential expression (DE) between GEX pixel clusters was performed by Wilcoxon rank sum test (Presto R package v1.0.0) using annotated genes. DE genes (adjusted p < 0.05, >20% detected expression in cluster) were visualized by heap map (ComplexHeatmap R package v2.12.1) using the top 10 overexpressed genes from each GEX cluster. Pseudotime trajectory analyses were done using the standard Monocle 3 pipeline (R package v1.3.1) with data that were already normalized using Seurat ^114^.

### Comparative analysis of pooled and perturb-DBiT CRISPR screen detection

Perturb-DBiT was benchmarked against the traditional pooled CRISPR screen detection using frozen liver sections from the Wang et al. screen of liver tumorigenesis genes ^40^. Briefly, a receiver-operator-curve (ROC) was constructed using sgRNA detection in the pooled screen data (>25% detection across tumor sample lobes) as the predictor for the perturb-DBiT sgRNA area (area > 2) as the response. The ROC and area-under-curve analyses were performed using the pROC R package v1.18.0.

### Preprocessing spatial screen data

A custom shell script was created to preprocess the screen data for perturb-DBiT using a combination of Cutadapt (v3.4) ^82^, Bowtie (v1.3.0) ^95^, and custom scripts. Briefly, a “whitelist” of possible unique molecular identifiers (UMIs) was created from the data using cutadapt with the following settings: -e 0.1 -m 10 -M 12 -l 10 --max-n 0 --discard-untrimmed. Next, alignment reference libraries were built for the UMI whitelist, as well as for the gRNA spacer sequences and the pixel barcodes using Bowtie-build. All barcodes were extracted from fastq files using cutadapt with the following:

| <u>Sequence type</u> | <u>Cutadapt parameters</u> |
| --- | --- |
| Spatial barcode A | -e 0.1 -m 8 -M 10 -l 8 --discard-untrimmed |
| Spatial barcode B | -e 0.1 -m 8 -M 10 -l 8 --discard-untrimmed |
| UMI | -e 0.1 -m 10 -M 12 -l 10 --discard-untrimmed |
| sgRNA spacer | -e 0.1 -m 15 -M 30 --discard-untrimmed |

The Cutadapt flanking regions were also specified by the perturb-DBiT chemistry:

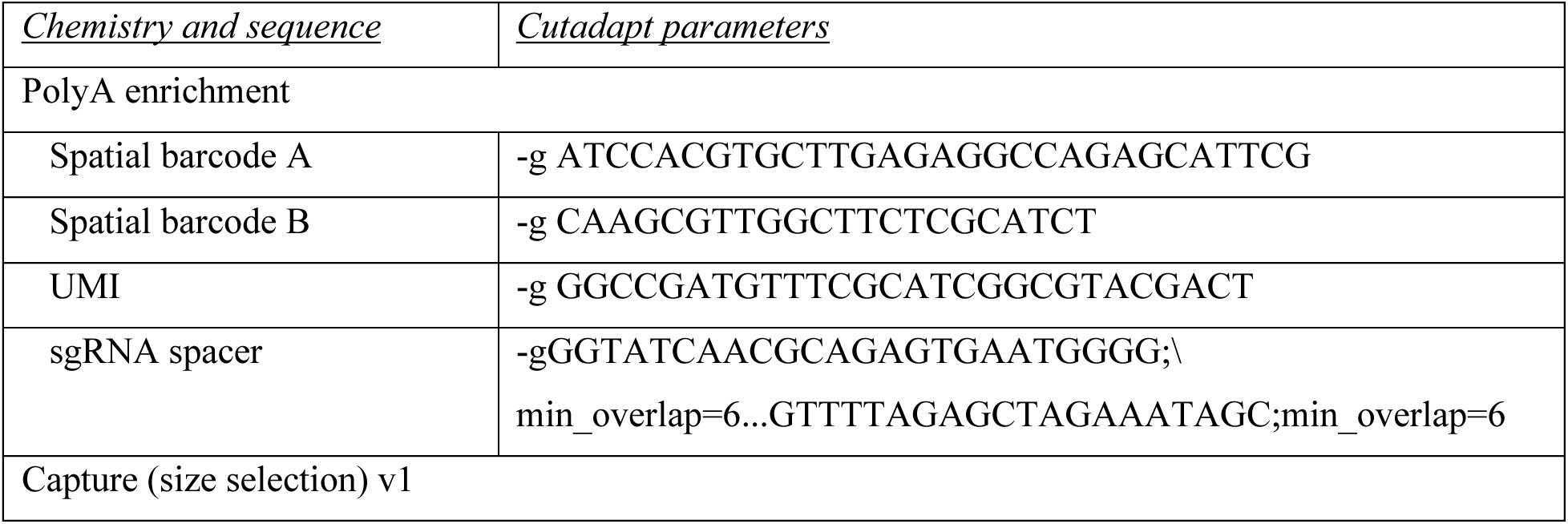
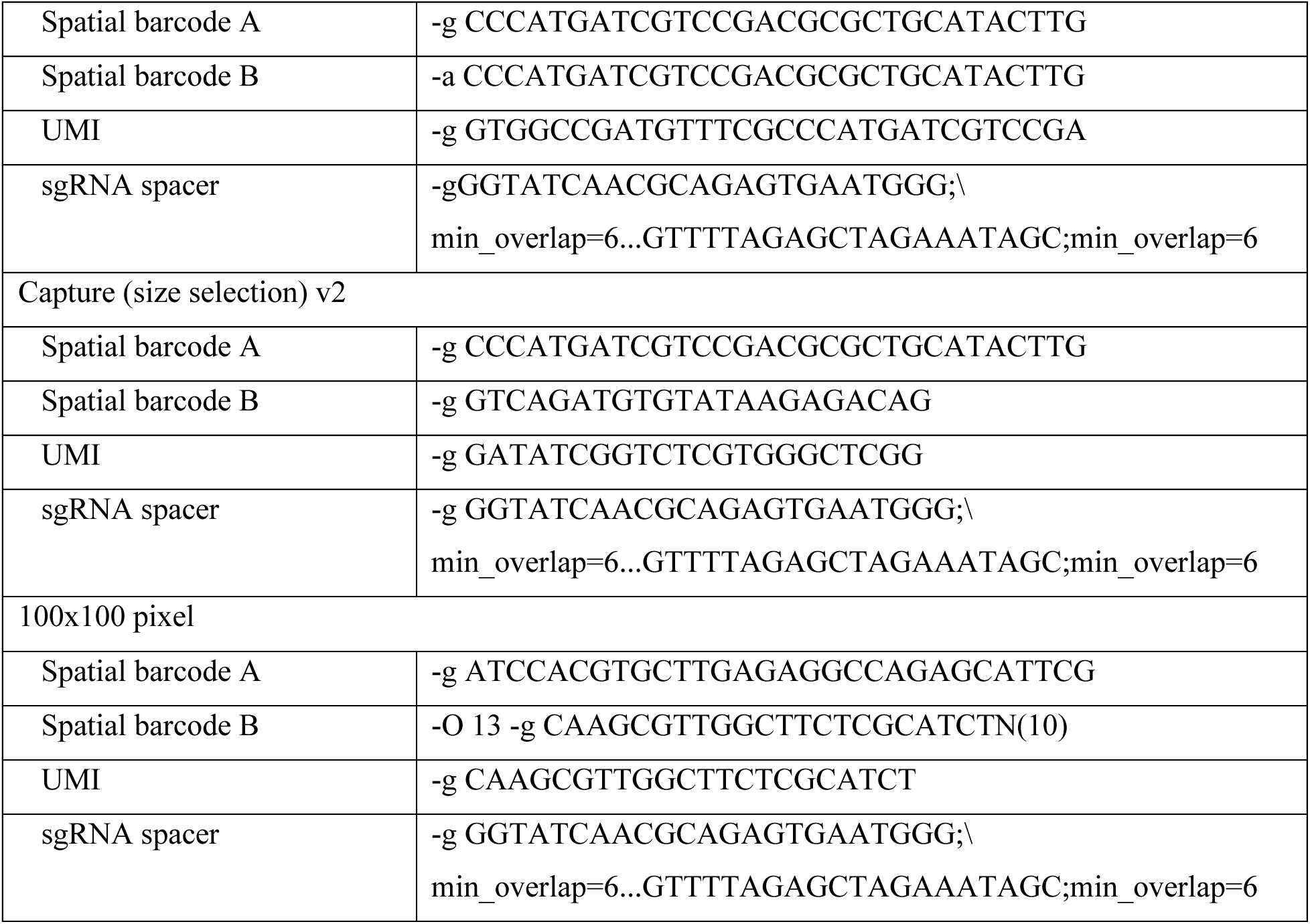

The trimmed sgRNA spacer sequences and barcodes for spatial position and UMI were then aligned to the custom libraries using Bowtie with the parameters below. Note that the parameters for mismatches and multi-matches (-v and -m, respectively), were optimized based on the reference library size and empirical testing with manually aligned reads.

| <u>Sequence type</u> | <u>Bowtie parameters</u> |
| --- | --- |
| Spatial barcode A | -v 1 -m 2 --suppress 2,4,5,6,7,8 --chunkmbs 2000 --best |
| Spatial barcode B | -v 1 -m 2 --suppress 2,4,5,6,7,8 --chunkmbs 2000 --best |
| UMI | -v 0 -m 4 --suppress 2,4,5,6,7,8 --chunkmbs 2000 --best |
| sgRNA spacer | -v 2 -m 2 --suppress 2,4,5,6,7,8 --chunkmbs 2000 --best |

A position ID was created from the aligned spatial barcodes, which were then matched to corresponding the corresponding sgRNA name and UMI for each paired sequencing read ID. Lastly, the sgRNAs were quantified based on the UMI count and subsequently analyzed with the R programming language.

### Perturb-DBiT screen analysis

Screen sgRNA quantification data were added to the GEX Seurat objects. The pixels were labeled by the uniquely detected sgRNA, or non-perturbed, if no sgRNAs were detected. Next, pixels with multiple sgRNAs were marked as ambiguous or resolved if the top sgRNA of the pixel had >= 10 counts and all other sgRNAs had 1 count. Genetic perturbation was analyzed using the Mixscape pipeline (Seurat R package v5.0.1) with slight modifications ^113^. First, perturbation signatures were calculated for all sgRNAs that were detected with an area >= 4 (all sgRNAs were used for samples with <100 different sgRNAs), and Mixscape analysis was performed, splitting data by cluster. The Mixscape results were filtered to get a list of perturbed pixels (perturbation-score > 0.2) and control pixels (NP pixels with perturbation-score < -0.2). All gene perturbation- scores for the filtered pixels were then re-arranged into a Seurat data assay. The perturbation assay was dimensionally reduced by PCA, and the top PCs (explaining the majority of the data variance) were further reduced by UMAP. Pixels were then clustered using the same strategy as described for GEX clustering.

Clustered pixels were labeled by the major perturbations of the cluster (perturbed gene for >20% of cluster pixels). Clusters with no major perturbations were labeled as “NP+n”, where n is the number of detected perturbations in the cluster. The perturbation clusters were characterized by the gene expression of liver tumor and metastasis-associated genes ^115–117^.

The gene expression pattern of each perturbation cluster was assessed by DE analyses using the NP perturbation cluster as the control and the same methods used for GEX DE. Pathway analyses were performed by Gene Set Enrichment Analysis (GSEA; fgsea R package v1.22.0), using biological process gene ontologies (2023 version, size <= 500 genes) ^104^, log2 fold-changes as the GSEA input, DE genes (adjusted p < 0.2), and the following parameters: function = fgseaMultilevel, gseaParam = 0.1, minSize = 2, and maxSize = 500. Lastly, the results were filtered to remove highly ambiguous pathways: “Translation”, “Transcription”, and “Gene Expression”.

### High-resolution tissue architecture analysis using iStar

The iStar algorithm is made up of three major components: a histology feature extractor, a high-resolution gene expression predictor, and a tissue architecture annotator, as detailed a previous publication^96^. H&E images were divided into smaller sections and then analyzed using a vision transformer to extract histological features. Then, a feed-forward weakly supervised trained neural network, then predicted superpixel-level gene expressions utilizing the top 1,000 most variable genes from the Perturb-DBiT expression matrix or top 16 sgRNA hits from the Perturb-DBiT sgRNA expression matrix. Clusters based on gene expression were used to segment the tissue and then were used for co-registration with the H&E-stained images. The top 10 genes associated with each cluster were identified and interpreted biologically, with input from a pathologist to refine the results based on tissue morphology.

### Co-registration and overlay between gRNA data and its histology

The brightfield image used in Perturb-DBiT was co-registered to its serial H&E-stained microscopy image via a manual control point based non-rigid co-registration workflow using a spline-based transformation model from Weave platform for Spatial Biology (Aspect Analytics NV, Genk, Belgium). The location of sgRNA measurements was precisely detected after co-registering its Barcode A and B images to the corresponding brightfield image. For this same section alignment, an automatic intensity matching co-registration workflow from Weave platform was used to find an affine transformation. After co-registering the brightfield image to its serial H&E-stained microscopy image via the manual non-rigid co-registration workflow, the sgRNA data was then precisely and directly linked to this same histology image. The pathologist annotated the H&E-stained microscopy image and placed 5 labels: ‘Bronchus’, ‘Vessel’, ‘Perivascular Fat’, ‘Peripheral Tumor’, and ‘Tumor Core’ via a web-based annotation tool included in the Weave platform. Finally, the sgRNA data and its serial histology data was precisely co-registered, overlaid and interactively visualized on the web-based Weave platform.

### Pseudotime analysis

Pseudotemporal analysis was performed with Monocle3^58^. Using scripts documented by a previous study^118^. Briefly, we normalized the count matrix with the preprocess_cds function (num_dim = 25). Dimensionality reduction was performed with the reduce_dimension function (reduction_method=’UMAP’, umap.min_dist = 0.5, preprocess_method=’PCA’). Next, we used reversed graph embedding with the learn_graph function to generate a principal graph from the reduced dimension space and we used the order_cells function to order the observations in a pseudotemporal manner. Clustering analysis was performed with the cluster_cells function at a resolution of 0.1 and the graph_test function was used to identify differentially expressed genes.

### Ligand-receptor interaction analysis

The R toolkit Connectome V1.0.0^103^ was utilized to analyze cell-cell connectivity patterns from ligand and receptor expression within our Perturb-DBiT dataset. The normalized Seurat object served as input, and UMAP clusters were used to define specific nodes within interaction networks. This resulted in an edge list that connected pairs of nodes based on their ligand-receptor mechanisms. Top-ranked interaction pairs were prioritized based on biological and statistical significance.

### Preprocessing of CODEX data

Cell segmentation was performed using a StarDist-based model in QuPath. The DAPI channel served as the nuclear marker for cell segmentation. The mean intensity of each marker per segmented cell was exported together with the centroids of each cell as a CSV. Downstream analysis was performed using Seurat 4.3.0 package. The dataset was normalized and scaled using the NormalizeData and ScaleData functions. Linear dimensional reduction was then performed with the “RunPCA” function. The “FindNeighbors” function embedded spots into a K-nearest neighbor graph structure based on Euclidean distance in PCA space, and the “FindClusters” function was used to cluster the spots. The “RunUMAP” function was used to visually show spatial heterogeneities through the Uniform Manifold Approximation and Projection (UMAP) algorithm. The clusters were plotted spatially using the ImageDimPlot function. The FindAllMarkers function was used to find the differentially expressed proteins in each cluster and the heatmap plotted using the DoHeatMap function.

### Statistical analysis

Prism V9 (GraphPad) was used for statistical analyses and specific tests are indicated in the text.

## Supplemental Table Titles and Legends

Table S1. Summary of samples and sgRNA data quality. Related to Figure 1.

Table S2. Lists of CODEX antibody list, cycle design, and reporter plate composition. Related to STAR Methods.

Table S3. List of primers and barcodes. Related to STAR Methods.

Table S4. List of sgRNAs and spacer sequences pertaining to Brie, Brunello, mTSG and Epigenetic libraries. Related to STAR Methods.

Table S5. List of Primers used in T7E1 assay. Related to STAR Methods.

