## Supplemental Materials for "Spatially Resolved *in vivo* CRISPR Screen Sequencing via Perturb-DBiT"

### Supplemental Figures

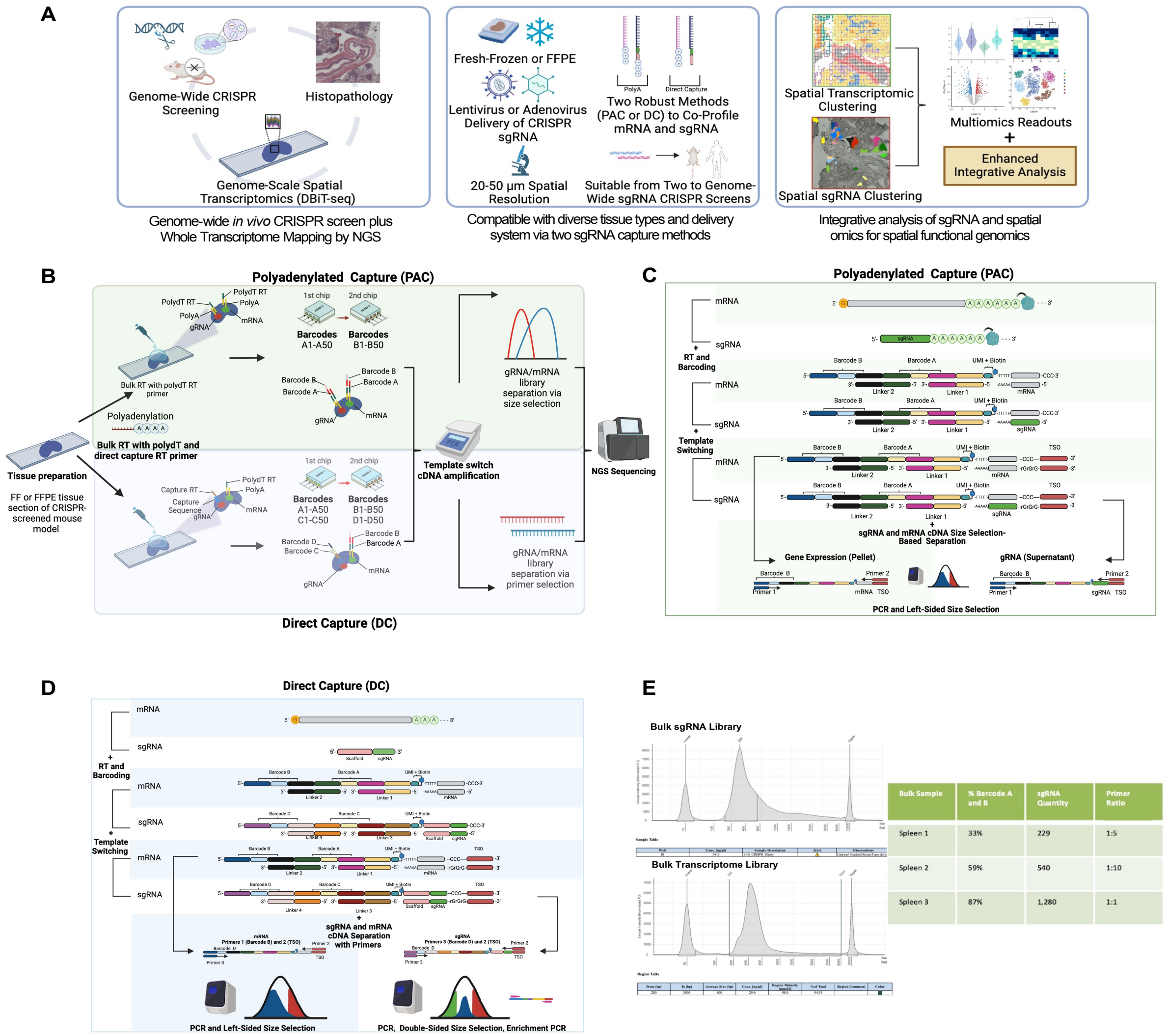


**Figure S1: Perturb-DBiT workflow, chemistry design, and optimization, Related to Figure 1**

**(A)** Schematic of Perturb-DBiT workflow PAC and DC methods
**(B)** Chemistry design of Perturb-DBiT PAC
**(C)** Chemistry design of Perturb-DBiT DC
**(D)** Optimization of Perturb-DBiT DC primer ratios via bulk experiment on fresh-frozen murine splenic tissue sections.

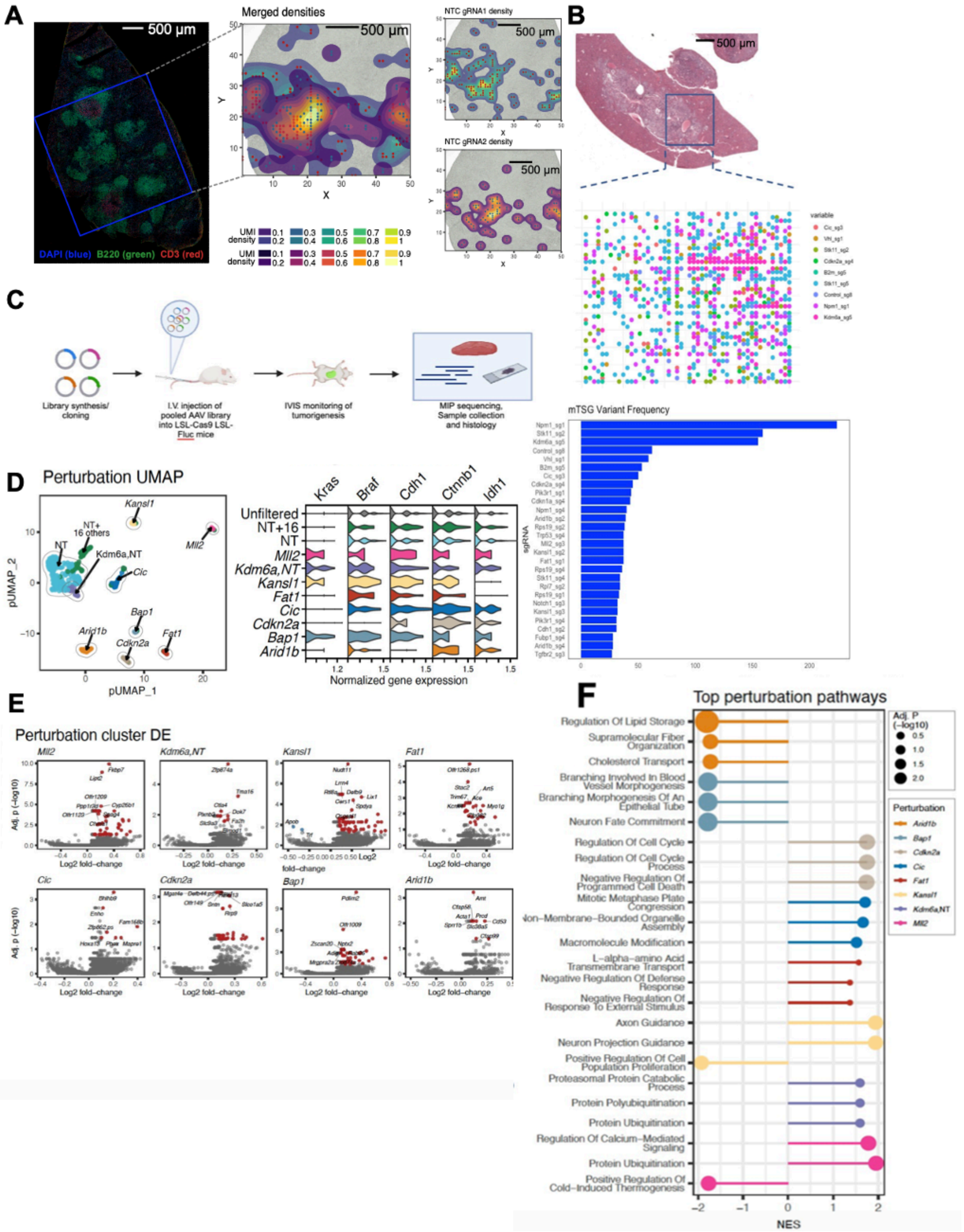


**Figure S2. Spatial mapping of sgRNAs in small and medium CRISPR-screening libraries, related to Figure 2**

**(A)** Small library guide detection (2 sgRNAs) in Cas9-expressing SMARTA cells in a mouse host spleen using a non-targeting control (NTC) dual-guide. Left, Immunofluorescence imaging of the host tissue is presented with host CD3+ T cells in red, and a blue box marks the ROI used for Perturb-DBiT. Right, The UMI detection of each of the dual guides are shown relative to the ROI. NTC sgRNAs are presented with points (sgRNA1 = red, sgRNA2 = blue) atop a 2D density map (sgRNA1 = blue-to-gold, sgRNA2 = purple-to-yellow).

**(B)** Top: Visualization of top sgRNA hits from 25-micron Perturb-DBiT detection of a medium-sized guide library (288 sgRNAs) in a murine model of autochthonous liver cancer. Bottom: Bar plot of top sgRNA hits from 25-micron Perturb-DBiT applied to a murine model of autochthonous liver cancer

**(C)** Schematics of direct *in vivo* AAV-CRISPR liver screen design.

**(D)** Left: UMAP visualization of pixels based on the dimensional reduction of Mixscape perturbation scores (pUMAP). Non-perturbed (NP) pixels were included in the analysis to serve as a control/reference point. Additionally, pixels were clustered (shown by color and outline) then named by the major perturbations of the cluster (>= 20% of clustered pixels). One cluster with NP and 16 minor perturbations (< 20% of cluster) was named "NP+16". Right: Violin plot of the expression of 5 selected oncogenes, compared between pUMAP clusters.

**(E)** Volcano plots of the DE analysis between selected pUMAP clusters using pseudo-replicate Wilcox tests. DE genes were those with an adjusted $p < 0.05$ and an absolute fold-change > 0.1 (up-regulation = red, down-regulation = blue), and the top 8 most significant DE genes were labeled for each analysis.

**(F)** Bubble plot of the top 3 pathways identified from DE genes of each pUMAP cluster (point color). Pathways were detected from gene set enrichment analyses using biological process gene ontologies, and the results are presented by significance (point size) and normalized enrichment score (NES), which provided the enrichment direction and magnitude.

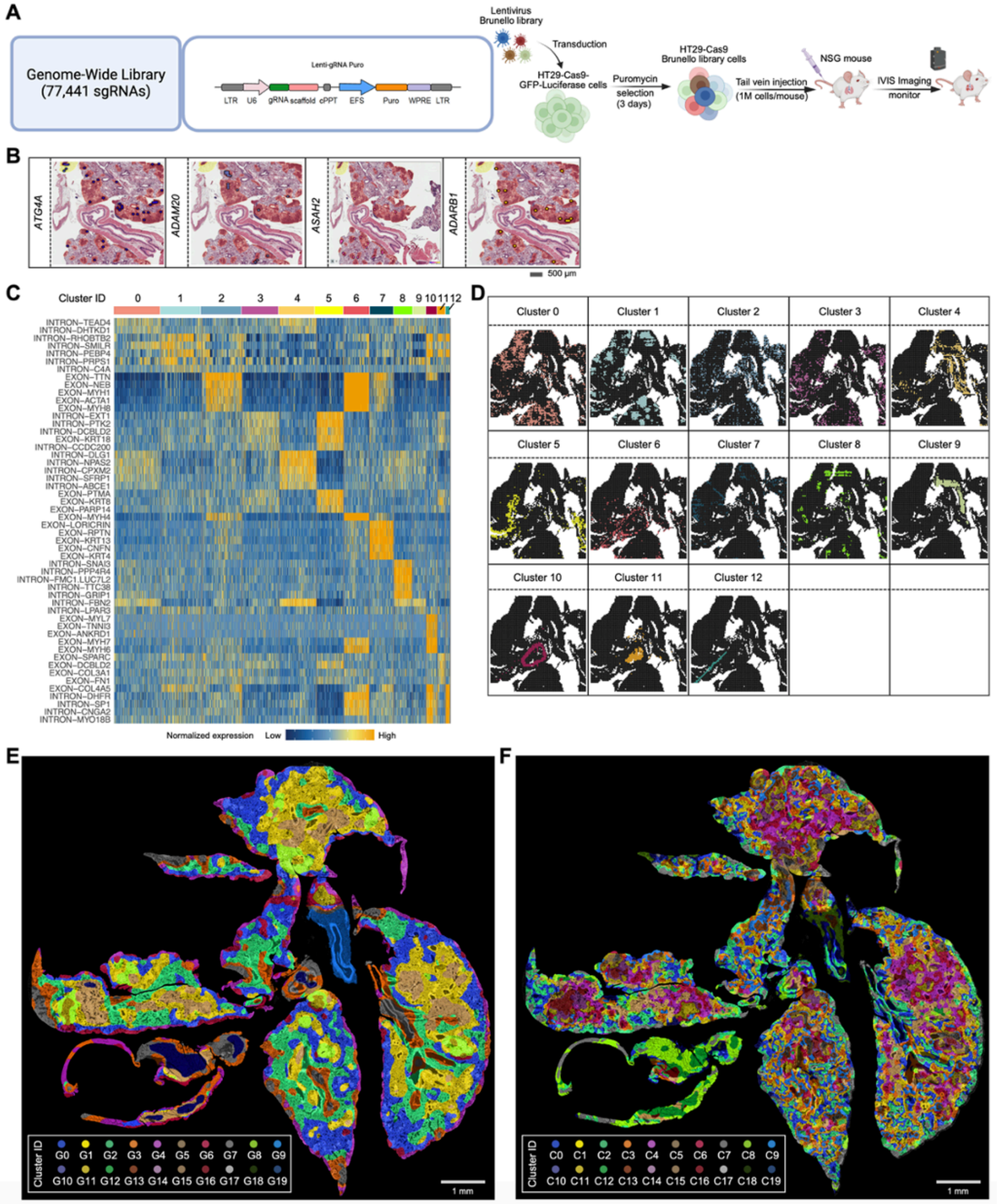


**Figure S3. Genome-wide high-resolution mapping of HT29 lung metastatic colonization model tissue sections, related to Figure 3**

(**A**) Left: Schematic of Lenti-gRNA Puro construct. Right: Schematic showing the development of HT29 lung metastatic colonization model.
(**B**) Spatial distribution of selected sgRNAs.
(**C**) Top ranked DEGs defining each cluster in Figure 3C.
(**D**) Spatial distribution of identified clusters.
(**E, F**) Super-resolved spatial clustering of the top 3,000 gene profile (E) or all sgRNAs (F) imputed with iStar.

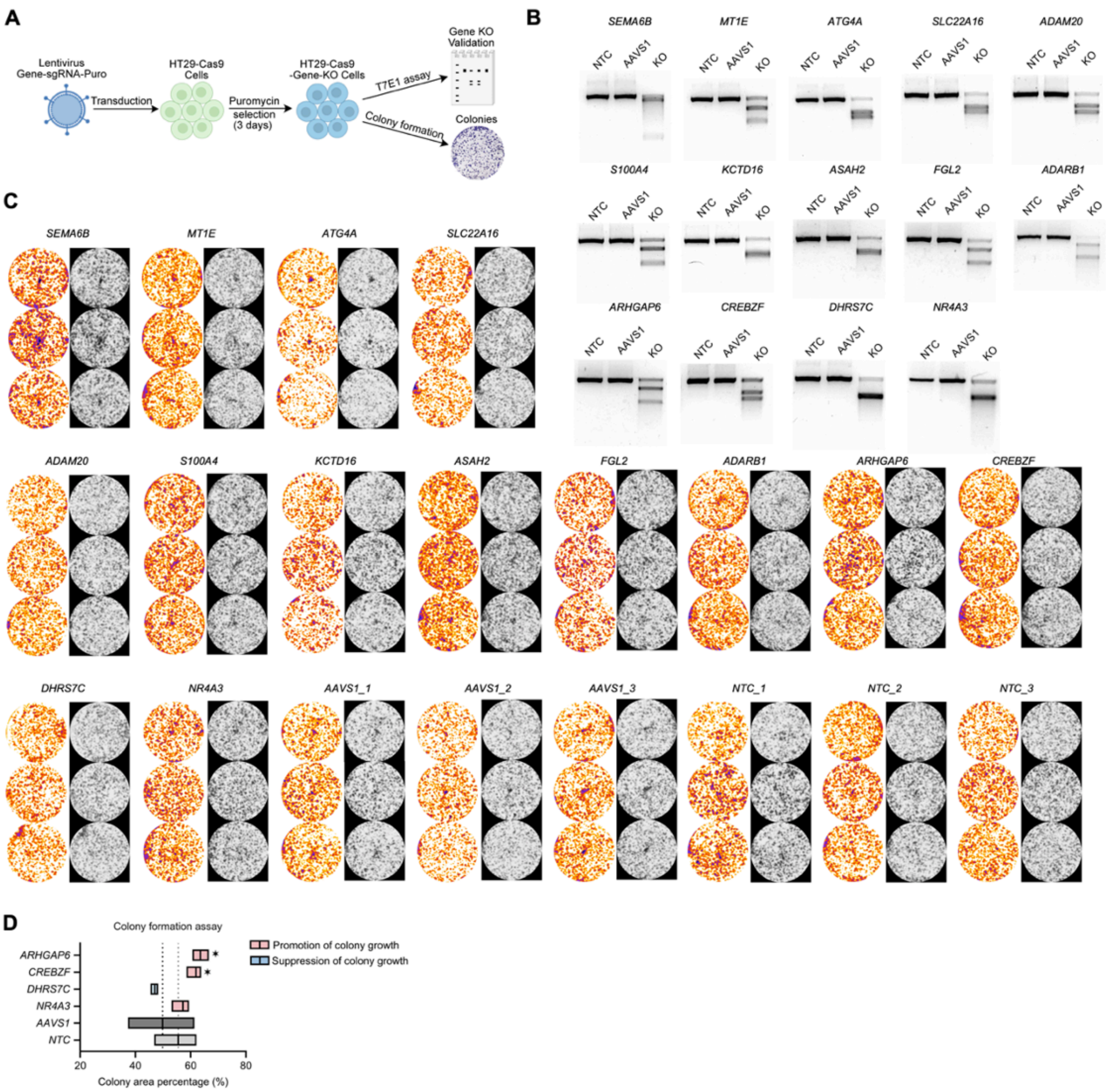


**Figure S4. Colony formation assays and spatial pseudotime analysis of HT29 lung metastatic colonization model, related to Figure 4**

(**A**) Schematic showing the colony formation assay.
**(B)** T7E1 assays determined genomic DNA editing and cutting efficiency of each sgRNA.
(**C**) Colony formation assay analysis utilizing Colony Area plugin in ImageJ of both top enriched sgRNA hits from Perturb-DBiT and validated tumor suppressor/promoter genes for the HT29 lung metastatic colonization model.
(**C**) Colony formation assay results of validated tumor suppressor/promoter genes for the HT29 lung metastatic colonization model.

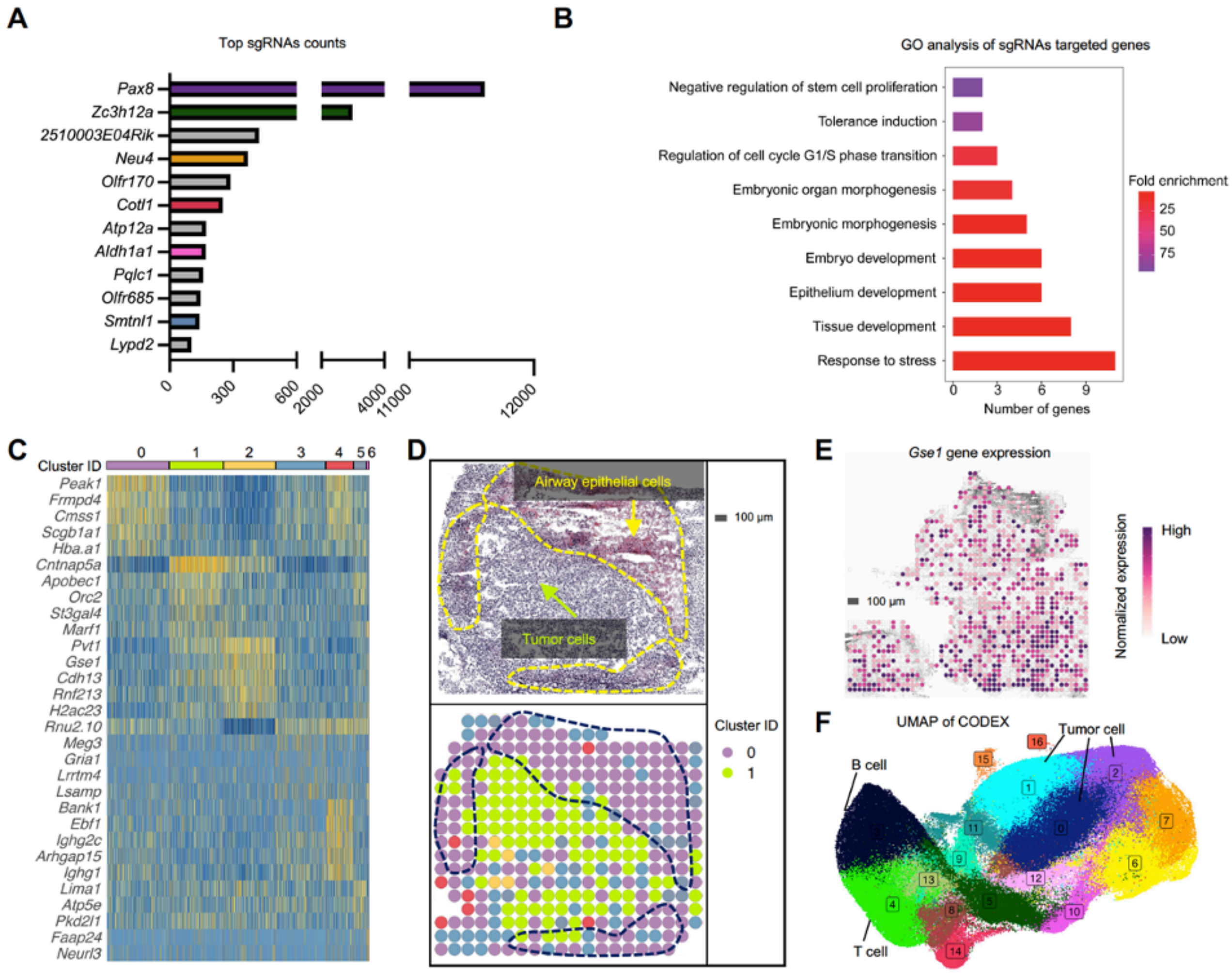


**Figure S5. E0771 syngeneic lung model schematic, sgRNA mapping and GO analysis, and tumor architecture insights, related to Figure 5**

**(A)** Schematic of E0771 syngeneic lung model.
**(B)** Spatial intensity plot of sample sgRNA detection via Perturb-DBiT.
**(C)** Top sgRNA hits and their respective counts detected by Perturb-DBiT.
**(D)** GO analysis of top sgRNA hits based on annotation from Ensembl and STRING-db.
**(E)** Heatmap of 6 distinct clusters revealed by unsupervised clustering of spatial transcriptomics data.
**(F)** Top: Pathology annotation differentiating airway epithelial cells and tumor cells within one of the three tumor regions covered by the ROI for Perturb-DBiT.
**(G)** Gene expression spatial plot of *Gse1* overlain on brightfield image of tissue section.
**(H)** CODEX UMAP revealing 16 unique clusters. Cluster 3: B cells, Cluster 4: T cells, Cluster 0,1,2: Tumor cells.
